# Maternal effects shape scale-dependent convergence in gut microbial responses to environmental change

**DOI:** 10.64898/2026.07.02.736155

**Authors:** Gabriele Schiro, Allyson Placko, Stan Boutin, Ben Dantzer, Jeffrey Lane, Andrew G. McAdam, Lauren Petrullo

**Affiliations:** Department of Ecology and Evolutionary Biology, University of Arizona, Tucson, AZ, USA; Biodesign Center for Health Through Microbiomes, Arizona State University, Tempe, AZ, USA; Department of Biology, University of North Carolina at Chapel Hill, Chapel Hill, NC, USA; Department of Biological Sciences, University of Alberta, Edmonton, Alberta, Canada; Department of Psychology, University of Michigan, Ann Arbor, MI, USA; Department of Ecology and Evolutionary Biology, University of Michigan, Ann Arbor, MI, USA; Department of Biology, University of Saskatchewan, Saskatoon, Saskatchewan, Canada; Department of Ecology and Evolutionary Biology, University of Colorado, Boulder, CO, USA

**Author notes:** **Corresponding author:** Gabriele Schiro.

## Abstract

Animals can experience environmental variability across multiple temporal scales, from predictable within-year seasonal fluctuations to rarer, irregular resource pulses that occur across generations. While gut microbiome responses to seasonal changes are well documented, far less is understood about how host-associated microbiomes respond to episodic environmental resource pulses, and whether such responses can persist across generations. Here, we investigate how gut microbial communities of wild North American red squirrels (*Tamiasciurus hudsonicus*) respond to intra-annual seasonal variation and episodic resource pulses happening years apart (specifically, seed masting). We then test the hypothesis that these responses are shared by mothers and their offspring. In line with prior work across mammals, we found that seasonal variation strongly structured gut microbiomes, driving widespread and convergent shifts in composition across the year. However, episodic resource pulses predicted microbial scatter such that gut microbiomes became more dissimilar across individuals in resource pulse years compared to typical years. Despite this population-level divergence, mothers and their adult offspring exhibited microbial convergence when encountering resource pulses occurring years apart, whereas seasonal responses were shared across the population and not unique to mother–offspring pairs. Overall, our results highlight the mammalian gut microbiome as a flexible component of the host phenotype, capable of adjusting to both predictable seasonal variation and large-scale, episodic resource pulses. Patterns of mother–offspring similarity across resource pulse years further provide evidence that some host-microbial responses to environmental change may be transmitted across generations.

## Introduction

Animals inhabiting fluctuating environments can experience ecological variability ranging from small-scale, regular seasonal changes to rarer, larger-scale ecosystem changes. For example, seasonality generates predictable changes in climatic conditions and resource availability that animals can track and rapidly respond to (Varpe, 2017). By contrast, rare or extreme disturbances, including droughts, wildfires, or storms, can dramatically reshape ecological conditions in an unpredictable manner. In between these two extremes are irregular but recurring events that animals may still sense and respond to, such as episodic phenomena like El Niño–Southern Oscillation (ENSO) dynamics (Marshal et al., 2011), or pulses of resources like prey items [e.g. cicadas or seed, (Yang, 2004)] and mast-seeding episodes (Clotfelter et al., 2007). Such variation across scales can challenge animals to maintain and optimize performance in dramatically different environments.

In mammals, gut microbiota may play a central role in mediating how animals adapt to these challenges, in part through shifts in microbial metabolic function that track changing environmental demands (Alberdi et al., 2016; Kolodny & Schulenburg, 2020). For example, seasonal environmental variation can trigger changes in host diet, immune function, or physiology that then reshape gut microbiota, driving further phenotypic change in the host (Bruce-Keller et al., 2015; Hooper et al., 2012; Lach et al., 2018). These microbiome-mediated changes can facilitate adaptation to fluctuating environmental conditions by enhancing host access to available resources and increasing physiological flexibility (Moeller & Sanders, 2020). For example, the gut microbiota can shift functional potential to more effectively degrade plant fiber or utilize host-produced mucus glycans as dietary availability changes across the year in gorillas (Hicks et al., 2018). Similarly, in wild wood mice, a transition from insect to seed based diet recapitulates seasonal shifts in gut microbial community structure (Maurice et al., 2015). Among hibernating animals, cyclical fasting shifts gut microbiota toward taxa capable of degrading host-derived substrates, in contrast to diet-driven changes during feeding periods (Carey & Assadi-Porter, 2017).

While studies continue to document seasonal host-microbiome dynamics, it remains unclear how gut microbial communities respond to ecological fluctuations that occur over longer inter-annual or multigenerational timescales. Similar to seasonal shifts, such changes can alter resource availability, diet, physiology and behavior; yet unlike seasonal shifts, episodic environmental fluctuations vary stochastically in their timing and magnitude, potentially generating more variable or idiosyncratic gut microbial responses across individuals and populations. One example comes from snowshoe hares, where gut microbial communities shifted not only across seasons but also became more diverse across inter-annual variation in population density during boom–bust cycles (Stothart et al., 2025), a pattern that could be only partially explained by dietary difference, suggesting that additional physiological mechanisms contribute to this restructuring.

To date, the evolutionary mechanisms by which microbial responses to environmental variation across seasonal to inter-annual timescales arise remain largely theoretical (Henry et al., 2021). The route through which hosts acquire their microbiome can influence not only microbiome composition but also how microbial communities respond and adapt to environmental challenges encountered by the host (Bruijning et al., 2022; Moeller & Sanders, 2020; Week et al., 2025). Bacteria can be incorporated into the gut microbiome from the surrounding environment (environmental transmission), from the individuals surrounding them [social transmission, (Moeller et al., 2018)], and through direct inheritance from parent to offspring during birth, nursing, or maternal care [vertical transmission, (Raulo et al., 2024; Valles-Colomer et al., 2023)]. These differences in transmission routes could have important evolutionary implications: microorganisms that depend on close contact for transmission may be more likely to evolve stable associations with their hosts (Leftwich et al., 2020) leading to potentially more mutualistic relationships (Moeller et al., 2018). A similar framework proposed by Bruijning et al., 2022 suggests that vertical transmission can favor microbial communities adapted to predictable, cyclical environmental changes, as vertically transmitted microbes may be shaped by natural selection to respond to environmental cues consistently encountered across generations.

Empirical evidence supporting these predictions remains limited in mammals. In sea anemones (*Nematostella vectensis)*, however, intergenerational transmission of heat-tolerant bacteria enhances tolerance to elevated temperatures, suggesting that vertical microbial transmission could represent a rapid mechanism of adaptation (Alberdi et al., 2016; Baldassarre et al., 2022). Alternatively, a more stochastic acquisition of microbiota from environmental sources when conditions vary unpredictably could confer an advantage by increasing functional flexibility, allowing hosts to integrate new microbes into their communities, and better respond to unforeseen environmental challenges. Weakened intergenerational microbial transmission could ultimately allow host populations to better hedge their bets when environments change unpredictably: for example, imprecise vertical transmission benefits marine sponge larvae facing variable environments (Björk, 2019).

In this study, we examine how mammalian gut microbiota vary across environmental fluctuations operating at multiple temporal scales, including regular within-year seasonal changes and episodic, predictable large-scale resource pulses. In the southwest Yukon, Canada, North American red squirrels (*Tamiasciurus hudsonicus)* experience regular, seasonal fluctuations in climate, food, and conspecific competition, as well as irregular environmental perturbations associated with their primary food source, seed from the cones of masting white spruce [*Picea glauca*, (Dantzer et al., 2020)]. Resource pulse or mast years, in which trees produce a superabundance of cones, occur only once every 3-7 years, with lower production of cones in the years in-between [non-mast years (Nienstaedt & Zasada, 2008)]. Spruce masting is the main driver of variation in individual short- and long-term fitness in this population (Hämäläinen et al., 2017), and squirrels have adapted to exploit these rare resource pulses by dramatically shifting behavior and reproduction in the months before the pulse occurs. Females adaptively increase reproductive rates ahead of the pulse by producing more offspring per litter as well as multiple litters [i.e., anticipatory reproduction, (Boutin et al., 2006; Dantzer et al., 2020; McAdam et al., 2019; Petrullo et al., 2023)]. Once the new cone crop is available in a mast year, red squirrels increase caching behaviors to collect orders of magnitude more cones (Fletcher et al., 2010). Both increases in breeding rates and cone collecting increase metabolic demands (Fletcher et al., 2012), and thus may be supported by metabolic reorganization of the gut microbiome during this time. Seasonal variation in gut microbiota across the year is known in this population (Ren et al., 2017), and red squirrel gut microbial variation can be shaped by social, behavioral, and physiological factors (Petrullo et al., 2022; Petrullo, Webber, et al., 2025). Nonetheless, the host-microbial response to resource pulses, its relative significance in shaping the gut microbiota compared to seasonal change, and the transmission mode that maintains it remains unknown.

Using over 700 fecal samples collected from 112 squirrels across 13 years (spanning 4 mast years), we first identified higher-order changes in patterns of gut microbial community composition with respect to seasonality and boom-bust changes in food availability. We identified the microbial taxa and functional pathways that demonstrated responsiveness to both smaller-scale, seasonal change and to large-scale, irregular changes in food availability associated with resource pulses. We hypothesized that recurrent environmental fluctuations at both scales favor the persistence and transmission of host-associated microbes across generations and predicted to find compositional convergence between mothers and their adult offspring sampled in similar seasons and years (resource pulse versus typical years).

## Methods

### Ethical statement

We obtained approval from the University of Michigan Institutional Animal Care and Use Committee, the University of Arizona Institutional Animal Care and Use Committee) and followed all relevant Canadian Council on Animal Care Guidelines and Policies. Fieldwork was permitted under Yukon Territorial Government Wildlife Research Permits and Scientists & Explorer’s Permits.

### Study system and population

All data for this study were collected across 13 years from a wild population of North American red squirrels (*Tamiasciurus hudsonicus*) living in the southwest Yukon, Canada. The Kluane Red Squirrel Project (KRSP) has continuously monitored red squirrels in this region since 1989 (Dantzer et al., 2020). Data were collected from squirrels living on one of two unmanipulated ∼40 ha study areas (“Kloo”/KL, “Sulphur”/SU) that experience similar environmental conditions. Monitoring consisted of biannual censuses of the entire population conducted in May and August/September. During each census, routine live trapping was conducted to record individual characteristics (e.g., body mass and pregnancy status) and collect fecal samples. Each spring, female squirrels were determined to be pregnant via behavioral observations of mating chases and abdominal palpation for the presence of fetuses. We outfitted pregnant females with radiocollars and used radio telemetry to locate nests within ∼24 hrs of females giving birth. At this time, pups were tagged for individual identification and parentage assignment.

### Fecal sample collection

We collected 735 fecal samples non-invasively from beneath traps during routine live-trapping from 112 squirrels (mean = 6.5 samples/individual, range = 1–33), across 13 different years between 2008 and 2023 (2013, 2020, and 2021 excluded), with 4 mast years (2010, 2014, 2019 and 2022). No juvenile individuals were included, as all individuals were born at least one year prior to sampling. Samples were stored on wet ice immediately after collection, transferred to a −20°C freezer within 5 h of collection, and kept at −20°C until further processing and long-term storage at -80°C at the University of Michigan and the University of Arizona. Samples were collected between day 41 and day 224 of the year (**supplementary information S1**). Samples were divided by season according to previous work (Ren et al., 2017): early spring: before day 120 of the year, late spring day 120 < 180 and summer > 180).

### Feeding observations

Feeding observations are collected opportunistically during fieldwork. That is registering what food items squirrels were consuming by visually identifying the item. In total, we recorded >27000 feeding events spanning the 13 years sampled (average 80 events per recorded week).

### DNA extraction, sequencing and bioinformatics

We extracted microbial DNA from fecal samples using the QIAamp PowerFecal Pro DNA Kit (Qiagen, Hilden, Germany) following manufacturer’s instructions. We amplified the V4 region of the 16S rRNA gene portion of the bacterial genome using primers 515F (5’-GTGCCAGCMGCCGCGGTAA-3’) and 806R (5’-GGACTACHVGGGTWTCTAAT-3’) following Caporaso et al. (2012). Primers contained Illumina adapter sequences, and reverse primers included unique 12-base pair Golay barcodes for demultiplexing. PCR amplicons were purified using QIAseq beads (Qiagen, Hilden, Germany) and quantified with the Quant-iT PicoGreen dsDNA Assay Kit (Invitrogen, Waltham, MA, USA). Purified products were pooled at equimolar concentrations and sequenced. Samples combined three different sequencing efforts, 324 and 251 samples were sequenced on two different runs on an Illumina MiSeq platform (Illumina, San Diego, CA, USA) using an 2x250 bp chemistry. An additional 139 samples were sequenced Illumina NextSeq 1000 platform (Illumina, San Diego, CA, USA), using an 2x150 bp chemistry. Sequences were resolved to Amplicon Sequence Variants (ASVs), which differ by only a single nucleotide, after initial quality filtering, denoising and chimera removal as implemented in the DADA2 pipeline [version 1.34; (Callahan et al., 2016)]. Taxonomy was assigned using the RDP classifier (Wang et al., 2007) as implemented in the DADA2 pipeline trained on silva v138.1 database (Quast et al., 2012). Extraction and PCR blanks were also sequenced to control for potential processing contamination. ASV tables from different runs were trimmed with the same parameters and merged after DADA2 using the function mergeSequenceTables as implemented in the DADA2 R package (Callahan et al., 2016). We generated predicted functional profiles for gut microbial communities using Phylogenetic Investigation of Communities by Reconstruction of Unobserved States (PICRUSt2) version v2.6.3. We obtained NSTI values (Mean = 0.12, Median = 0.12, quartiles 0.06–0.22), indicating similar if not better prediction precision compared to other mammalian microbiomes (Langille et al., 2013). Run-to-run differences were accounted for in all statistical tests performed in the study to control for potential batch effects.

### Community diversity metrics

All statistics were performed in R 4.2.1 (Team R Development Core, 2020). We first visualized alpha and beta diversity analysis on rarefied data (10000 reads). We assessed the effects of season and masting on microbial richness and Shannon H’ using linear mixed-effects models [lme4 package in R (Bates et al., 2007)]. In the case of Shannon H’, a power transformation was applied to achieve linearity. The model included season (early spring: before day 120 of the year, late spring day 120 < 180 and summer > 180), masting status, and their interaction as fixed effects, along with host age, sex, grid location, and sequencing read number as covariates. To account for non-independence in the data, we included random intercepts for individual squirrel ID, year, and sequencing run. We performed post-hoc pairwise comparisons between seasons and mast using estimated marginal means [emmeans package (Lenth & Piaskowski, 2025)]. We adjusted all p-values for multiple comparisons using Tukey’s Honest Significant Difference method.

For beta diversity, we constructed a Bray-Curtis distance matrix using vegan v.2.6.8 (Oksanen et al., 2018). We then ran a PERMANOVA including sequencing run, number of reads, sex, grid, age, year (as categorical), and season as covariates, with a season×mast (yes or no) interaction term. We used a conservative sequential testing approach (Type I SS) with technical and demographic factors ordered first, to be sure that season and mast effects represented variance beyond that explained by potential confounders. To account for repeated sampling of individuals, permutations were constrained within squirrel IDs (strata argument), controlling for individual-level variation while testing temporal and environmental effects (N=999 permutations).

### Differential enrichment and depletion of microbial taxa across environments

To identify genera that varied in relative abundance across seasons and masting conditions, we performed differential abundance testing on the 50 most abundant genera in the dataset. For each genus, we fit a generalized linear mixed model with a negative binomial distribution using the glmmTMB package in R. The model included season and mast year as fixed effects with their interaction term, and random intercepts for individual squirrel, year, and sequencing run. Total read count per sample was included as a log-transformed offset term. For each genus, we estimated marginal means across seasons and mast vs. non-mast conditions within each season using the emmeans package, and performed pairwise contrasts to test for differences between masting conditions. We adjusted p-values for multiple testing across all contrasts using the Benjamini-Hochberg false discovery rate (FDR) correction, with significance set at FDR < 0.05.

### Bayesian multimembership regression models

To test how gut microbial communities changed across seasons and between mast and non-mast years, we first calculated how similar each pair of samples was using Bray–Curtis and Jaccard similarity metrics. We then used Bayesian beta regression models in brms (Bürkner, 2017), to test which factors explained variation in microbiome similarity between pairs of samples. We tested community similarity differences between “mast matched” samples (e.g., samples that both come from mast years) and all other pairs of samples (e.g., mast/non-mast and non-mast/non-mast). We also compared samples of mother-offspring microbiome similarity between masting and non-masting years. As covariates in the model, we controlled for seasonal timing (same season vs. different), spatial distance, sex concordance, age differences, and technical factors (sequencing depth and run). To control for non-independence of the data structure, we included sample pairs and individual squirrel pairs as multiple membership structures [mm(); (Bürkner, 2017)]. Absence of collinearity between predictors was assessed calculating variance inflation factors. All predictors models had VIF < 3. Models were threaded across 4 parallel chains with 3,000 iterations each (1,000 warmup iterations). Convergence was confirmed with R-hat values < 1.01 and effective sample sizes > 400. We assessed statistical significance of fixed effects by examining whether credible intervals (95%) excluded zero on the logit scale. For visualization, we transformed parameter estimates from the logit scale to the probability scale using the inverse-logit transformation (plogis) and reporting the change in similarity with the baseline plogis(intercept + coefficient) - plogis(intercept). Post-hoc pairwise comparisons for interaction effects were conducted using estimated marginal means (emmeans package). To identify which bacterial genera drive mother-offspring microbiome similarity, we fit a separate ordered beta regression model (Kubinec, 2023) for similarities calculated within members of the 50 most abundant genera (ordered beta regression accounts for zero and one in the data, given that at genus level zero and one similarities between communities are common). For each genus, posterior estimates and 95% credible intervals on the response scale were extracted using the R library marginal effect (Arel-Bundock et al., 2024). We retained only genera with credible intervals not overlapping zero and a minimum effect size of |estimate| > 0.05. Distance in years was removed, as the single-genus model showed a strong correlation with the same year in the majority of cases.

## Results

### Red squirrel gut microbial communities are dominated by Firmicutes and Bacteroides

Across all gut microbial communities in our dataset, bacterial taxa belonged to 8,216 unique ASVs. At the phylum level, most ASVs belonged to Firmicutes (54.29% mean relative abundance), Bacteroidota (36.12%) and Cyanobacteria (4.09 %). The most abundant bacterial families were Lachnospiraceae (29.6 %), followed by Prevotellaceae (17.02 %) and Muribaculaceae (12.54 %) (**Fig 1A**).

**Figure 1.**
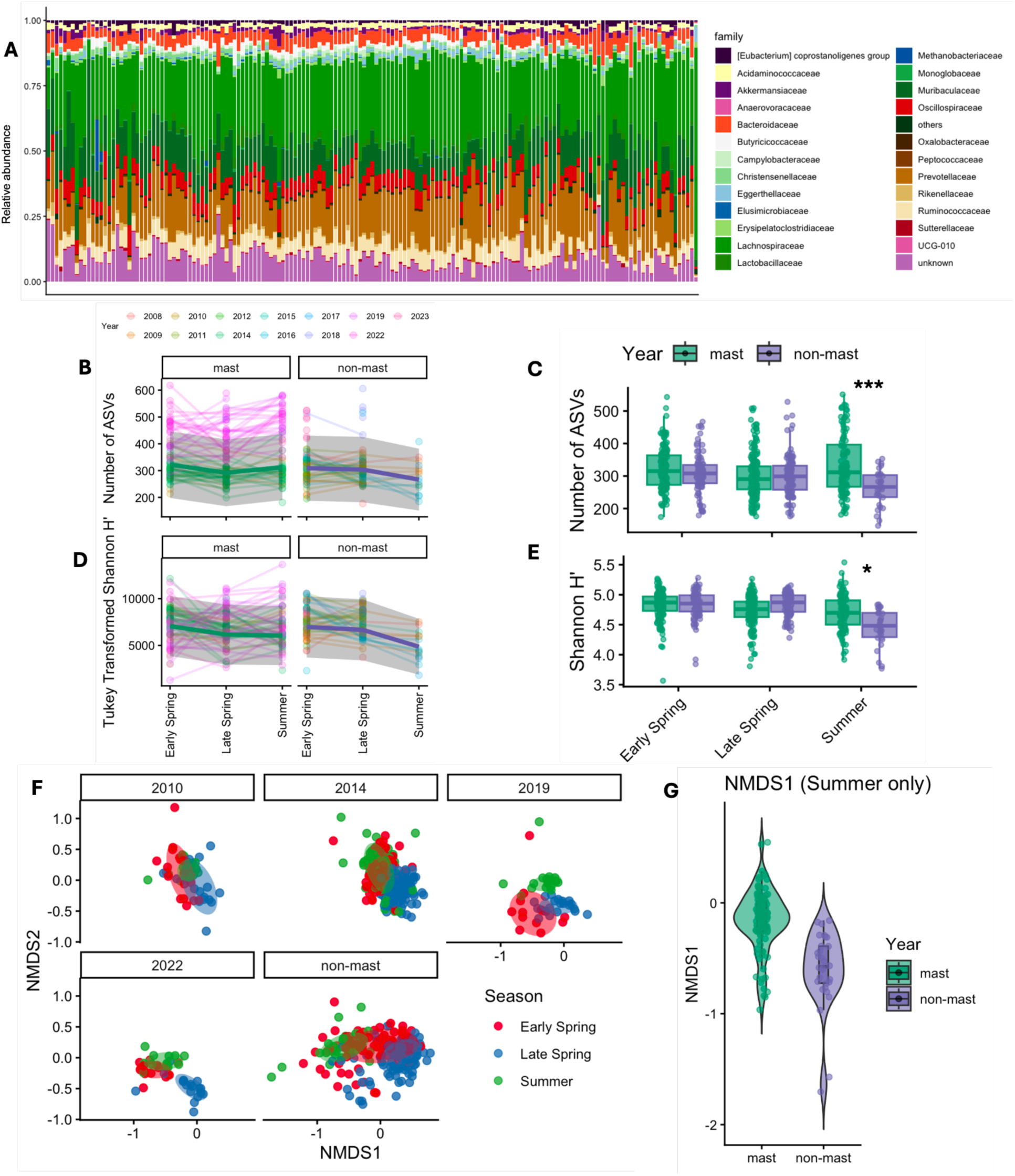
Gut microbiome structure reflects seasonal dynamics and resource pulses. (A) Taxonomic composition at family level, highlighting the most abundant bacterial clades in the dataset. Only the 25 most abundant families across the whole dataset are shown with others grouped under “others”. Bars are ordered according to the Julian calendar. (B) ASV richness and (D) Shannon diversity index (H’), across seasons in mast and non-mast years. Each colored line represents an individual squirrel across different years; the thick line is the predicted trend and gray shading showing 95% confidence intervals as modelled with a generalized linear mixed model. Seasonal differences in (C) ASV richness and (E) Shannon diversity index (H’), in mast and non-mast years. Results show lower diversity during summer in mast years. (F) NMDS ordination of microbial community composition colored by season for mast years (2010, 2014, 2019, 2022) and pooled non-mast years. Communities cluster by season, with a clearer separation of summer samples during mast years. (G) Violin plot showing differences in the primary ordination axis (NMDS1) between mast and non-mast years in summer samples.

### Individual microbial diversity changes across environmental scales

To test the effect of seasonality and resource pulses on individual microbial diversity, we modelled microbial richness (i.e., number of ASVs detected in each sample) and Shannon H’ diversity (abundance-weighted metric) using linear mixed effect models (tables with full results in **supplementary information S2 and S3**). Pairwise tests from these models (n = 679) demonstrated greater ASV richness (**Fig 1B**) in early spring than in either late spring (estimated difference = 17.7, P < 0.001) or summer (estimated difference = 25.9, P < 0.001). The effect of a resource pulse on microbial richness was not consistent throughout the year, but instead varied as a function of season (season × mast interaction: *F*_2, 586.05_ = 20.31, P < 0.001). In the spring, microbial richness remained at similar levels in mast and non-mast years, across both early and late spring periods (P > 0.05, **Fig 1C**). However, this pattern shifted in summer, with a strong increase in microbial richness in resource pulse years compared to typical years (estimated difference = 46.4, P < 0.001, **Fig 1C**). Measures of abundance-weighted gut microbial diversity demonstrated similar patterns, with greater diversity overall in early spring compared to late spring (Shannon H’ estimated difference = 611, P < 0.001, **Fig 1D**) and summer (Shannon H’ estimated difference = 1,569, P < 0.001, **Fig 1D**). Similarly, resource pulses increased abundance-weighted diversity only in summer (season × mast: F_2, 533_ = 7.24, *p* < 0.001; Shannon H’ estimated difference = 1,188, P = 0.045, **Fig 1E**).

### Resource pulses cause population-level shifts in microbial composition

Compositional variation in microbial communities (expressed as Bray-Curtis dissimilarity) demonstrated that community composition was shaped by different spatio-temporal factors (all predictors had P < 0.001, **supplementary information S4**). Year had the strongest effect on community variation (8.6% explained variation), while seasonal differences were also evident (5.1 % explained variation, **Fig 1F**). As resource pulses occur only within specific years, its independent effect could not be separated from year; nevertheless, it influenced how communities changed across seasons, with seasonal patterns differing between resource pulse and typical years (season × mast interaction 0.08%, **Fig 1F-G, supplementary information S4**).

To identify which bacteria were driving these differences, we quantified microbial taxa that were differentially enriched or depleted across seasons and resource pulse environments. 48 of the 50 most abundant genera consistently varied across seasons (**Fig 2A**). This included *Prevotellaceae.UCG.001*, the most abundant genus in the dataset, which was at its lowest in early spring and reached its peak during late spring regardless of mast conditions (**Fig 2B**). In mast years, significant shifts in the relative abundance of bacterial genera occurred mainly in summer, with thirteen genera exhibiting enrichment during this period in resource pulse compared to typical (non-mast) conditions (**Fig 2A**). Notably, some bacterial genera involved in the digestion of plant fibers strongly increased in the summers of mast years (e.g., *Ruminococcus;* mast year summer vs. non-mast year summer, estimated log_2_ fold-change 3.33, P_adj_ < 0.001, **Fig 2C**), while others involved in cellulose degradation decreased in mast year summers (e.g., *Marvinbryantia;* Summer Mast vs Non-Mast, estimated log_2_ fold-change-1.45, P_adj_ < 0.001, **Fig 2D**). Because both seasonal transition and mast years are associated with a shift in dietary intakes (see **supplementary information S5** for a dietary overview), these changes in observed taxa might reflect corresponding shifts in food resources. To evaluate this, we ran a mantel’s test between diet and microbiome composition, which showed a positive correlation between the two distance matrices, indicating that such changes can be attributed to diet (mantel’s spearman rho = 0.24, P < 0.001).

**Figure 2.**
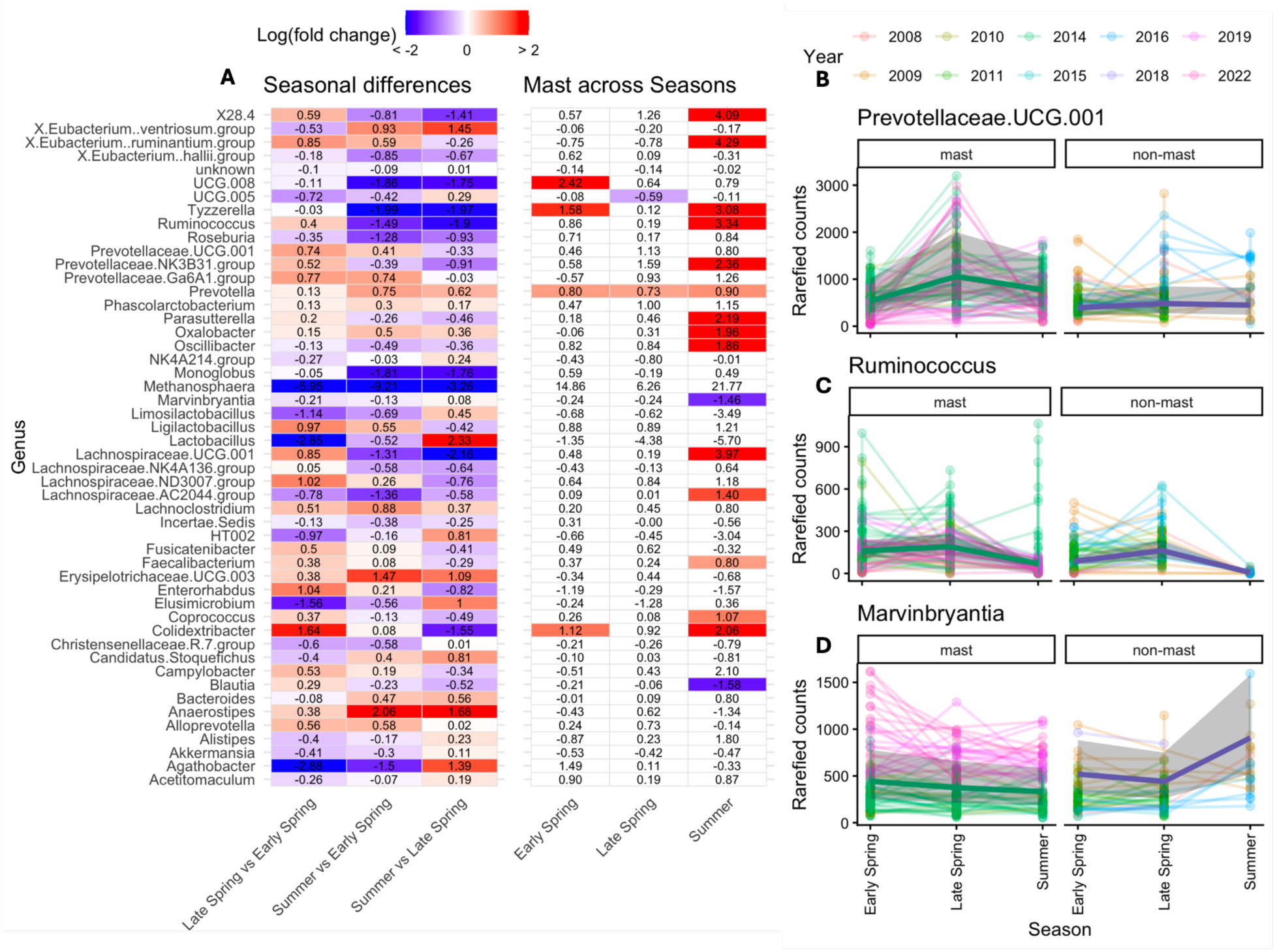
A subset of bacteria become differentially enriched across seasons and resource pulse years. (A) Heatmap showing changes in abundance, expressed as log(fold change) in bacterial genera abundances across seasons (left three columns) and between mast and non-mast years within each season (right three columns). Only genera with significant differential abundance are colored (p-adj < 0.05 after FDR correction). (B-D) Temporal patterns across seasons for three representative genera: (B) *Prevotellaceae* UCG-001, the most abundant genus across the whole dataset, show pronounced differences across seasons. (C) *Ruminococcus*, showing both seasonal trends but also a significant increase in summers of mast years, and (D) *Marvinbryantia*, which instead, decreases. Line plots show rarefied genus abundance trajectories from early spring through summer, separated by mast (left) and non-mast (right) years. Colored lines represent individuals across different years, and the black line is the predicted trend and gray shading showing 95% confidence intervals as modelled with a generalized linear mixed model.

### Both seasonality and resource pulses shape microbial metabolic potential

Seasonal comparisons revealed a pronounced shift in functional pathway profiles, with later seasons, particularly summer, showing a clear enrichment of metabolic pathways relative to spring periods (late spring vs early spring 151 pathways significantly increasing and 83 decreasing, summer vs early spring 352 increasing and 30 decreasing, summer vs late spring 248 increasing and 51 decreasing, after P_adj_ < 0.05, **Fig 3A**). These changes were spread across different functional categories (**Fig 3A**). The predominance of enriched pathways in summer suggests community wide shifts in microbial functional potential and increased functional convergence. Microbial communities become more similar in their metabolic profiles over time, resulting in a greater number of pathways reaching statistical significance (full results of post hoc comparisons in **supplementary information S6**).

**Figure 3.**
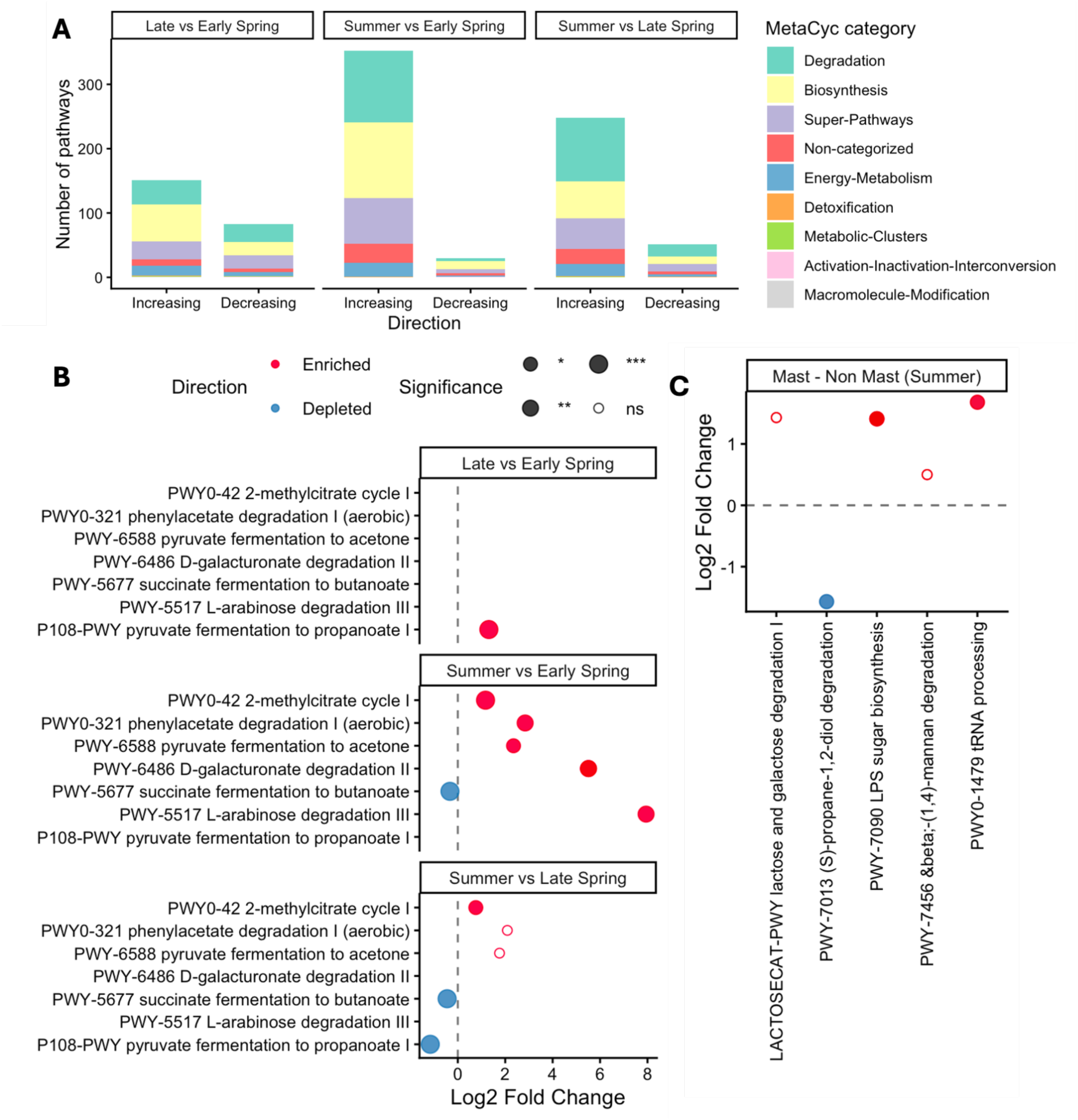
Gut microbiome metabolic potential declines toward summer and shows enhanced biosynthetic capacity with resource pulses. (A) Number of significantly changing MetaCyc pathways across seasonal transitions (early spring–late spring, early spring–summer, late spring–summer), grouped by functional category. Pathways are classified as increasing or decreasing based on the direction of log-fold change- (FDR < 0.05), revealing an increase of pathways in summer, indicating seasonal expansion of metabolic potential across multiple functional categories. (B) Log (fold changes) of selected pathways involved in fermentation and compound degradation across seasonal comparisons. These results highlight contrasting trends, with enriched pathways related to intermediate metabolite processing in summer (e.g., methylcitrate cycle, phenylacetate degradation) and depletion of pathways associated with plant-derived carbohydrate degradation (e.g., D-galacturonate and L-arabinose degradation), indicating a shift in substrate utilization across seasons. (C) Differential abundance of metabolic pathways between food boom and non-boom years during summer, the only season showing significant differences. Positive log₂ fold-changes indicate enrichment during food boom years. Pathways related to biosynthesis and RNA processing are enriched under food boom conditions, whereas select degradation pathways are reduced, suggesting altered metabolic activity in response to increased resource availability.

Several pathways related to fermentation and intermediate compound processing shifted across seasons. For example, pyruvate fermentation to propanoate (P108-PWY) peaked in late spring (**Fig 3B**), while pyruvate fermentation to acetone (PWY-6588) and 2-methylcitrate cycle (PWY0-042) were enriched in summer (**Fig 3B**). In contrast, succinate fermentation to butanoate (PWY-5677) was depleted in summer compared to early spring (**Fig 3B**), suggesting a reduction in butanoate-producing fermentation pathways. These results indicate a shift in dominant fermentation end-products across the year. Pathways involved in the degradation of plant-derived substrates were also differentially abundant across seasons. Notably, L-arabinose degradation III (PWY-5517), D-galacturonate degradation II (PWY-6486) and phenylacetate degradation I (aerobic) (PWY0-321) were enriched in summer relative to early and late spring (**Fig 3B**), indicating increased utilization of hemicellulose- and pectin-derived sugars and enhanced breakdown of aromatic compounds towards summer. Comparisons between diet and microbiome community functional potential revealed a weak but significant positive correlation between the two distance matrices, indicating that temporal changes in functional potential were linked to changes in diet (Mantel Spearman’s rho = 0.20, P < 0.001). Of the 535 pathways that were differentially enriched or depleted across seasons, 187 also demonstrated significant positive correlations with dietary items (P_adj_ < 0.05, **supplementary information S7**). For example, new cones, which are consumed predominantly during summer mast years, were associated with the broadest functional response and increased functional diversity spanning multiple metabolic classes (**supplementary Information S8**). By contrast, spruce bud consumption was primarily associated with polysaccharide and polyol degradation pathways, particularly inositol metabolism, a compound present in conifer buds (Pullman et al., 2008). Stored cones elicited a functional response in the gut microbiota that was distinct from that elicited by new cones, suggesting that storage alters microbial processing of seeds and elicits a different gut microbial response (**supplementary information S8**).

Across boom-bust environments, only three pathways were significantly differentially abundant (P_adj_ < 0.05, **Fig 3C**). Specifically, PWY-7090 Lipopolysaccharide (LPS) biosynthesis and PWY0-1479 tRNA/nucleotide processing pathways were enriched in mast years (PWY-7090 estimated log_2_ fold-change 1.41, P_adj_ = 0.01, PWY0-1479 estimated log_2_ fold-change 1.68, P_adj_ = 0.02), suggesting enhanced bacterial protein translation and bacterial growth (**Fig 3C)**, while a decrease in PWY-7013 (*S*)-propane-1,2-diol degradation pathway (estimated log_2_ fold-change - 1.57, P_adj_ = 0.02) suggests changes in fermentation strategies of the community. When a less conservative (10%) significance threshold was used, we found summer increases in LACTOSECAT-PWY (indicating in lactose and galactose degradation potential, estimated log_2_ fold-change 1.43, P_adj_ = 0.07) and PWY-7456 (indicating increased mannan degradation potential,estimated log_2_ fold-change 0.5, P_adj_ = 0.07), consistent with increased breakdown of seed-derived carbohydrates (Downie & Bewley, 2000). When compared to dietary intake, PWY-7090, PWY0-1479 and PWY-7456 were significantly correlated with stored (but not fresh) cone consumption (PWY-7090: Spearman’s rho = 0.25, P_adj_ = 0.02; PWY0-1479: Spearman’s rho = 0.3, P_adj_ = 0.005; PWY-7456 Spearman’s rho = 0.33, P_adj_ = 0.002, **supplementary information S7**).

### Mother–offspring pairs converge in response to large but not small-scale environmental changes

To test whether parent–offspring transmission of microbiota (i.e., vertical transmission) influences how squirrel gut microbiomes change across seasons and population boom–bust cycles, we used pairwise Bayesian multi-membership models to compare the gut microbiomes of pairs of squirrels. These models test whether pairs of squirrels that are more similar in their traits (e.g. same sex, sampling year, spatial proximity, etc), also have more similar microbial communities, while accounting for the paired and non-independent structure of the data. Full model outputs on logit scale are presented in **supplementary information S9-10**. We constructed pairwise Bayesian multimembership models using both Jaccard (presence/absence of microbial taxa) and Bray–Curtis (community distribution of taxa) and metrics of microbial similarity (Raulo et al., 2024). Pairs of squirrels that were sampled in the same year (Jaccard estimate 0.09, CI 0.08 0.09; Bray-Curtis estimate 0.12, CI 0.11 0.12) and were of the same sex (Jaccard estimate 0.003, CI 0.001 0.005; Bray-Curtis estimate 0.006, CI 0.003 0.009) had more similar gut microbiomes. Pairs of squirrels with greater temporal (distance in years between samples, Jaccard estimate -0.24, CI -0.25 -0.24; Bray-Curtis estimate -0.26, CI -027 -0.25) and spatial (distance in meters between samples, Jaccard estimate -0.09, CI -0.10 -0.09, Bray-Curtis estimate -0.11 CI -0.12 -0.10) distance between them exhibited more dissimilar gut microbial communities by both metrics (**Fig 4A-B**).

**Figure 4.**
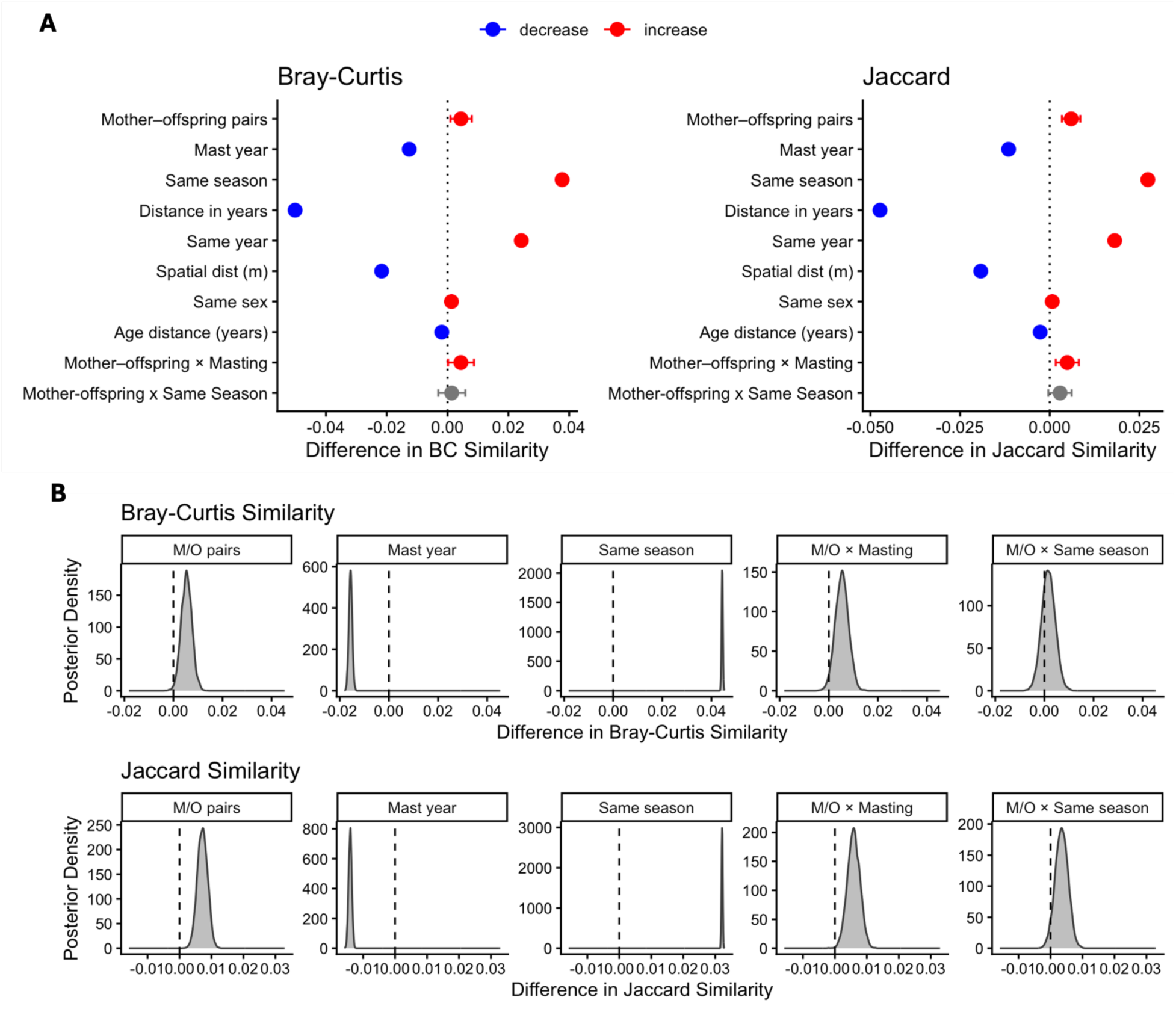
Host relatedness and environmental factors shape gut microbiome similarity across individuals. (A - B) Effects of host-related and environmental predictors on pairwise microbiome similarity based on Bray–Curtis (A) and Jaccard (B) dissimilarities, and (C), posterior distributions of key model effects for Bray–Curtis (top) and Jaccard (bottom) similarity. Points represent posterior mean estimates from Bayesian multimembership models, with error bars indicating uncertainty. For both metrics, mother–offspring pairs show increased similarity, consistent with vertical transmission. Shared environmental conditions (same season, same year) also tend to increase similarity, whereas spatial distance and temporal distance reduce similarity. Notably, **f**ood boom years decrease similarity in indicating divergence in composition. In contrast, the interaction between mother–offspring pairs and food booms is positive, suggesting increased convergence among related individuals during resource pulses despite broader community divergence.

Environmental changes exhibited divergent effects on pairwise microbial similarity depending on scale. Seasonal changes predicted broad gut microbial convergence, with pairs of squirrels sampled in the same season (even if in different years) exhibiting significantly more similar gut microbial communities in terms of both the presence/absence of specific microbes (Jaccard estimate: 0.13, CI: 0.12 0.13) and community distribution of those taxa (Bray-Curtis estimate: 0.18, CI: 0.18 0.18, **Fig 4A-B**). By contrast, food boom environments caused strong differentiation between microbial communities, suggesting that large-scale environmental disturbances trigger individual-specific microbial responses. Pairs of squirrels both sampled in mast years (including different mast years) had less similar microbial communities compared to pairs sampled in non-mast years [Jaccard estimate: -0.06, CI: 0.06 0.05; Bray-Curtis estimate: - 0.06, CI: -0.07 -0.06, **Fig 4A-B**)].

These temporal dynamics were modified by parent-offspring microbial transmission. In general, mother-offspring pairs exhibited greater microbial similarity across both metrics compared to unrelated/other pairs, indicative of vertical microbial transmission [Jaccard estimate: 0.03, CI: 0.02 0.04; Bray-Curtis estimate: 0.02, CI: 0.01 0.03, (Ren et al., 2017), **Fig 4A-B**)]. Food boom environments further magnified this effect, such that the interaction between mother-offspring pairs and masting was positive for both presence-absence (Jaccard estimate: 0.02, CI: 0.01 0.04, **Fig 4A-B**) and community structure (Bray-Curtis estimate: 0.02, CI: 0.001 -0.04, **Fig 4A-B**) measures of microbial similarity. In other words, mast years amplified mother-offspring convergence in gut microbiome composition, indicating that mothers and their offspring exhibit similar microbial responses to food booms even when those booms occur in different years. By contrast, season only weakly predicted mother-offspring convergence, and did not reach statistical significance [Jaccard estimate: 0.01, CI: 0.08 0.09, Bray-Curtis estimate: 0.007, CI: - 0.001 -0.03, **Fig 4A-B**)].

### *Prevotellaceae* are shared across generations and respond to environmental change

To identify the bacterial genera that underlie patterns of microbial similarity across seasons and resource pulses, we estimated genus-level effects using Bayesian regression models for both presence–absence and community distribution measures (**Fig 5**). Increased spatial distance between samples was associated with reduced microbial similarity in both measures among genera *Alloprevotella, Prevotellaceae Ga6A1 Prevotellaceae UCG001* (all estimates < -0.05 and CIs not crossing zeros). Samples collected in the same year also exhibited convergence across nine different genera for Bray-Curtis (all estimates > -0.05 and CIs not crossing zeros, **Fig 5**), and five for Jaccard (all estimates > -0.05 and CIs not crossing zeros, **Fig 5**). Genera *Colidexteribacter*, *Roseburia* and *UCG.005* (Oscillospiraceae) increased their Jaccard similarity in samples coming from matching seasons, potentially indicating seasonally specific acquisition patterns (all estimates > 0.05 and CIs not crossing zeros for same season), with other two genera showing only season specific Bray-Curtis convergence (*Anaerostipes* and *Lachospiraceae NK4A136 group*) pointing to season specific responses (all estimates > 0.05 and CIs not crossing zeros for same season, **Fig 5**).

**Figure 5.**
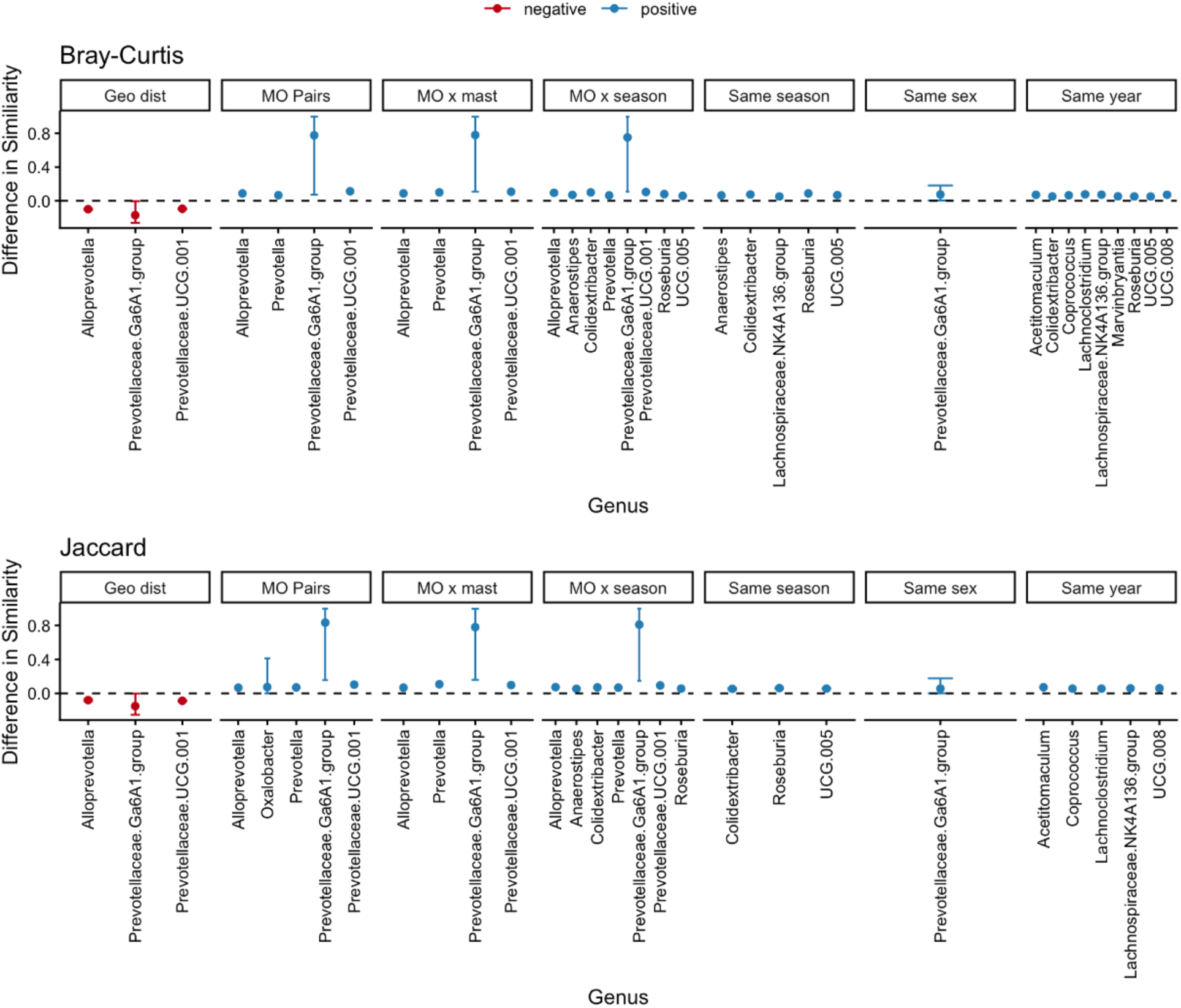
Genus-specific responses to host relatedness, seasonality, and temporal distance. *Prevotellaceae* genera are primarily structured by host-related factors and shared environmental conditions, whereas other genera are more strongly influenced by temporal distance and interannual variation. Each point represents the posterior mean estimate (± 95% credible intervals) from Bayesian regression models (*brms*) of the effect of a given predictor on pairwise microbiome similarity for a given bacterial genus, with only statistically meaningful effects shown (credible intervals not overlapping zero and |estimate| > 0.05). Positive values indicate increased similarity, whereas negative values indicate reduced similarity. Across both Bray–Curtis (top) and Jaccard (bottom) metrics, mother–offspring pairs and their interactions with food booms and season consistently increase similarity, particularly for genera belonging to *Prevotellaceae*. In contrast, geographic and temporal distance (year difference) reduce similarity, with stronger effects observed for non-*Prevotellaceae* taxa, indicating greater temporal turnover in these groups.

Across both measures, mother–offspring microbial convergence was consistently predicted by members of the bacterial family Prevotellaceae (*Alloprevotella, Prevotella, Prevotellaceae Ga6A1,* and *Prevotellaceae UCG001*, all estimates > 0.05 and CIs excluding zeros, for both Bray-Curtis and Jaccard similarity, mother offspring pairs), the genus *Prevotellaceae Ga6A1* group had the strongest estimate for this comparison (Jaccard estimate -0.06, CI 0.06 0.05; Bray-Curtis estimate -0.06, CI -0.07 -0.06, **Fig 5**). The mother-offspring convergence exhibited by these taxa was further strengthened in resource pulse years across both metrics of microbial similarity (*Alloprevotella, Prevotella, Prevotellaceae Ga6A1,* and *Prevotellaceae UCG001*, all estimates > 0.05). Interestingly, convergence between mother–offspring pairs also increased within the same season for these genera, while additional genera (*Colidexteribater, Roseburia* and *Anaerostipes)* showing a similar trend (all estimates > 0.05 and CIs not crossing zeros, for both Bray-Curtis and Jaccard similarity, mother offspring pairs mast interaction **Fig 5**). This indicates that part of the responses to seasonal changes can also be transmitted from mother to offspring; however, the weaker overall community signal suggests a more widespread response across the host population.

## Discussion

Investigating how host-associated microbial communities shift across temporal and environmental gradients is fundamental to understanding their role in shaping host metabolic flexibility and, ultimately, the capacity of hosts to cope with changing conditions (Alberdi et al., 2016). Overall, we found that within-year seasonal variation shaped gut microbial community composition more strongly than across-year resource pulses. Seasonal changes during summer months predicted a sharp decrease in individual microbial richness and abundance-weighted diversity, and strong enrichment of microbial functional pathways related to metabolism. This pattern suggests homogenization of the functional capacity of the community, where broad functional diversity is lost in favor of bacteria that perform a narrower but perhaps more ecologically or physiologically relevant set of functions.

Specifically, seasonal shifts in fermentation pathways (P108-PWY, PWY-6588, PWY0-042, PWY-5677; propanoate, acetone, methylcitrate, and butanoate production) and plant-derived substrate degradation (PWY-5517, PWY-6486, PWY0-321; arabinose, galacturonate, and aromatic compound breakdown), point to both possible differences in food inputs and energetic demands regardless of resource pulse conditions. Consistent with this interpretation, we found a positive correlation between temporal changes in microbiome functional potential and dietary composition, and several of these pathways were also significantly associated with diet, further supporting a link between seasonal dietary shifts and microbiome functional restructuring. Food items were associated with distinct functional signatures, with buds linked primarily to sugar and polysaccharide degradation pathways, stored cones to growth-related biosynthetic pathways, and new cones to the broadest range of metabolic functions. These results are consistent with previous studies in this population that link compositional shifts in the gut microbial communities with shifts in feeding behavior and diet (Ren et al., 2017). Additionally, territorial behaviors and fluctuations in population densities can homogenize microbiomes at the population level and diversify microbiomes at the individual level, potentially through social microbial transmission (Petrullo, Webber, et al., 2025). Because territorial dynamics vary seasonally (Webber et al., 2023), these behavioral changes may also contribute to some of the observed seasonal differences in gut microbiome composition demonstrated here.

Although seasonal effects were generally stronger, we also captured significant effects of resource pulses on gut microbial community composition. Differences between regular and of resource pulse years were most pronounced in the summers, coinciding with the period in which squirrels are increasing reproduction and fresh new cones are available. We specifically detected changes in the relative abundance of a number of key fermentative bacteria. For example, the bacterial genera *Ruminococcus* and *Lachnopspiraceae UCG001* were enriched and *Blautia* and *Marvinbryantia* were depleted. All four of these genera contribute to fiber digestion and degradation in their hosts (Esquivel-Elizondo et al., 2017; Hartinger et al., 2022; Rusling et al., 2024)*. Ruminococcus* also has been observed to promote increased movement and activity in wild mice, potentially through production of short-chain fatty acids, and by enhancing glucose release from resistant starches (Rusling et al., 2024). Higher abundance of *Ruminococcus* sp. CAG 382 were associated with increased litter size in sows, suggesting a potential contribution to enhanced reproductive performance through serotonin-related host–microbiome interactions (Chen et al., 2025). Mast year enrichment of *Ruminococcus* may therefore support the increased energetic demands associated with collecting new cones and sustaining higher reproductive outputs characteristic of mast years summers (Boutin et al., 2006; Fletcher et al., 2012). We also observed a depletion of functional pathways associated with 1,2-propanediol degradation. 1,2-propanediol is an intermediate produced during microbial fermentation of dietary fibers and host-derived substrates and is commonly involved in cross-feeding among gut microbes (Cheng et al., 2020). Finally, our finding of enrichment in microbial pathways related to the degradation of mannans and galactose, both found in spruce seed, point to an increased ability to digest seed in resource pulse years (Downie & Bewley, 2000).

This concentration of changes during the period of resource emergence suggests that microbiome shifts are likely responses to resource changes and/or the energetic demands associated with these changes rather than anticipatory adjustments ahead of a disturbance (Petrullo, Morris, et al., 2025). However, some microbial taxa exhibited enrichment before the period of new cone availability in resource pulse years, pointing to anticipatory shifts in community ahead of a large-scale environmental shift. In particular, bacteria in the genus *Prevotella* became significantly enriched in early spring and remained enriched through summer in mast years, with no significant enrichment during parallel periods in non-mast years. *Prevotella* are commonly present in rodent microbiomes and are involved in fermentation and the release of short-chain fatty acids that can serve as metabolic signaling molecules for hosts (Gruninger et al., 2016; Kohl et al., 2014). In voles, *Prevotella* are involved in the regulation of reproductive hormone synthesis (Zhu et al., 2022), suggesting that anticipatory enrichment of *Prevotella* in resource pulse environments may be involved in the anticipatory reproductive and behavioral strategies that squirrels use to maximize fitness in mast years [e.g., increased reproduction, (Boutin et al., 2006; McAdam et al., 2019; Petrullo et al., 2023)].

At the population level, pairwise analyses revealed that resource pulses reduced microbial similarity across both microbial presence-absence (i.e., Jaccard dissimilarity) and abundance-weighted (i.e., Bray-Curtis dissimilarity) metrics, leading to population-wide divergence in gut microbiomes. Given that we found no bacterial genera that similarity increased in masting years, these findings point to a “masting signal” that causes shifts across different functional groups, and indicates that microbial turnover occurs primarily across genera (or through abundance shifts between genera), rather than within genera. This pattern of microbial scatter could be caused by individual differences in behavioral responses to resource pulses: for example, reproductive investment generally increases in response to cues of an upcoming spruce mast (Boutin et al., 2006; Hämäläinen et al., 2017; Petrullo et al., 2023), but individuals can vary in the degree of this investment (Hämäläinen et al., 2017; Petrullo et al., 2023). It may also be caused by individual differences in diet in resource pulse years if some squirrels incorporate the new seed that is produced in the summer months into their diets earlier, or at greater magnitudes, than others. More broadly, diet-associated functional analyses showed that consumption of new cones in general was associated with the broadest range of microbial metabolic functions, supporting increased functional diversification during periods of high new resource availability. Furthermore, the reduced similarity among individuals could also be explained by the Anna Karenina effect, whereby host stress caused by increased energetic demands in mast years could weaken the physiological and immune mechanisms that normally regulate gut microbial communities, allowing stochastic processes of community assembly to cause divergent microbiome compositions (Ma, 2020; Zaneveld et al., 2017).

In mammals and some other taxa, vertical microbial transmission homogenizes mother-offspring gut microbial communities through direct transfer of maternal microbiota to the offspring gut during and after birth (Asnicar et al., 2017; Björk, 2019; Bruijning et al., 2022). Here, we demonstrate significant convergence in the gut microbiota of mothers and their offspring, consistent with vertical transmission (Ren et al., 2017). Interestingly, this mother–offspring convergence disrupted the pairwise differentiation of gut microbial communities predicted by food booms. Mast years exacerbated dissimilarity in gut microbial communities among individuals except for between mother-offspring pairs, which instead exhibited microbial convergence across different resource pulse environments. This reversal suggests that, while large-scale ecological disturbances may cause microbial scatter broadly, the microbial response to such disturbances may have an inherited component. This inheritance would be consistent with theoretical predictions that vertically transmitted microbes could be particularly important in changing but predictable environments (Bruijning et al., 2022). It is also consistent with the assumption that vertical transmission, as a mechanism of host-microbe co-diversification, can promote the maintenance of microbiota that not only fill vital functional needs for their hosts (Leftwich et al., 2020; Murphy et al., 2023), but that aid hosts in responding to fitness-relevant challenges consistently experienced across generations (Bruijning et al., 2022; Leftwich et al., 2020). As a deterministic mechanism, vertical microbial transmission may underlie the adaptive phenotypic plasticity that is favored in populations inhabiting variable but predictable environments (Levins, 1968). While convergence during mast years may reflect the amplification of inherited microbial lineages that are adapted to food boom cues, some carryover differences across offspring could still act as a source of population-level variation buffering against random extinction events (Murphy et al., 2023).

Microbial convergence among mother–offspring pairs in mast years may also emerge when offspring mirror the physiological and behavioral responses of their mothers. For example, intergenerational similarities in territorial and reproductive behaviors during mast years could lead to parallel shifts in microbiome composition through similar microbial exposures and/or host physiological demands. Behavioral and reproductive traits such as aggression, activity, and litter size in this population possess low but significant heritability, as well as substantial maternal effects (Taylor et al., 2012). If offspring inherit tendencies in territoriality or reproductive investment from their mothers, they may exhibit similar dietary shifts and behavioral changes in response to cues of an upcoming mast. Territory bequeathal is also a recurring behavior in this population (Berteaux & Boutin, 2000), allowing some offspring to remain on or near the natal territory, increasing environmental overlap between mothers and offspring and potentially further augmenting mother-offspring microbiome similarity.

At the taxonomic level, bacteria belonging to the family *Prevotellaceae* demonstrated a consistent increase in mother–offspring similarity across both abundance-weighted (Bray–Curtis) and presence-absence (Jaccard) metrics, suggesting a persistently high degree of vertical transmission from mother to offspring among members of this genus. *Prevotella* was also the genus that exhibited increasing abundance during mast years throughout all studied seasons. Its concurrent association with mast years and strong mother–offspring similarity raises the possibility that microbiome responses to resource pulses are maintained across generations through the vertical transmission of this taxon. Specifically, bacteria within the genus *Prevotellaceae* Ga6A1 exhibited the strongest effect. In a previous study on the same study system, periods of heightened social density negatively predicted the relative abundance of *Prevotellaceae* Ga6A1 across individual gut microbiome communities (Petrullo, Webber, et al., 2025), suggesting vertical rather than social routes of acquisition for this particular genus.

We expected to similarly find mother-offspring microbial convergence in response to seasonal change. However, mother-offspring pairs did not exhibit detectable microbial convergence when sampled in the same season. This may indicate weaker selective pressures for the vertical transmission of microbiota that respond to seasonal variation, perhaps because individuals can acquire the microbes needed to cope with seasonal changes from their broader environment, or because individuals can rely on a more generalized and less specific set of microbes to meet seasonal demands [i.e., functional redundancy among various microbial taxa (Moya & Ferrer, 2016)]. We note that this does not indicate that vertically transmitted microbes do not respond to seasonal change. At the genus level, some seasonal responses may still be maintained by vertical transmission. In particular, increases in similarity in these genera were detected for the interaction between season and mother–offspring pairs. Instead, it suggests that the response of gut microbiota to seasonality may be widespread across the population and largely independent of vertical transmission and/or maternal effects. The weaker signal at the community level suggests that, while certain taxa exhibit maternally linked seasonal dynamics, the dominant pattern remains a population-wide response rather than one driven by maternal transmission.

Together, our findings provide evidence that while large-scale environmental disturbances like resource pulses can shape microbial responses, some microbial shifts can occur in advance of the pulse, pointing to a capacity for anticipatory microbial reorganization alongside anticipatory host phenotypic changes. That mother-offspring gut microbial communities converged across resource pulse years but not in response to seasonal change, despite resource pulses otherwise generating microbial differentiation, suggests that vertical transmission may mechanistically promote the maintenance of microbiota with the capacity to respond to cues of large-scale environmental changes. This may be most likely to emerge in host populations that have evolved to track and predict these changes. Ultimately, these data highlight the gut microbiome as a flexible part of the host phenotype, capable of adjusting to both predictable seasonal changes and episodic resource pulses, with potential intergenerational inheritance of microbial responses to large-scale disturbances.

## Availability of data and materials

Sequencing raw files are available on the NCBI sequencing read archive with the access code PRJNA1478286. Code is available at https://github.com/Petrullo-Lab/squirrel_mast_project.

## Acknowledgements

We thank the Champagne and Aishihik First Nations for allowing us to conduct our work within their traditional territory. We thank all field technicians who have contributed to data collection over the years. We thank Dr. Aura Raulo for valuable discussions on the Bayesian model setup. Sequencing was performed at the University of Michigan Microbiome Core and the University of Arizona PANDA Core for Genomics and Microbiome Research. Bioinformatic analyses were conducted using High-Performance Computing (HPC) resources supported by the University of Arizona. This work was supported in part by the Natural Sciences and Engineering Research Council of Canada (S.B., J.E.L., A.G.M) and the National Science Foundation (A.G.M., IOS-1749627 to B.D.).

## Author contributions

G.S. and L.P. conceived the study and wrote the manuscript. A.P. conducted lab work. All authors contributed edits and revisions to all versions of the manuscript.

## References

1. Alberdi, A., Aizpurua, O., Bohmann, K., Zepeda-Mendoza, M. L., & Gilbert, M. T. P. (2016). Do Vertebrate Gut Metagenomes Confer Rapid Ecological Adaptation? Trends in Ecology & Evolution, 31(9), 689–699. 10.1016/j.tree.2016.06.008

2. Arel-Bundock, V., Greifer, N., & Heiss, A. (2024). How to Interpret Statistical Models Using **marginaleffects** for *R* and *Python*. Journal of Statistical Software, 111(9). 10.18637/jss.v111.i09

3. Asnicar, F., Manara, S., Zolfo, M., Truong, D. T., Scholz, M., Armanini, F., Ferretti, P., Gorfer, V., Pedrotti, A., Tett, A., & Segata, N. (2017). Studying Vertical Microbiome Transmission from Mothers to Infants by Strain-Level Metagenomic Profiling. mSystems, 2(1), e00164–16. 10.1128/mSystems.00164-16

4. Baldassarre, L., Ying, H., Reitzel, A. M., Franzenburg, S., & Fraune, S. (2022). Microbiota mediated plasticity promotes thermal adaptation in the sea anemone Nematostella vectensis. Nature Communications, 13(1), 3804. 10.1038/s41467-022-31350-z

5. Bates, D., Sarkar, D., Bates, M. D., & Matrix, L. (2007). The lme4 package. R Package Version, 2(1), 74.

6. Berteaux, D., & Boutin, S. (2000). BREEDING DISPERSAL IN FEMALE NORTH AMERICAN RED SQUIRRELS. Ecology, 81(5), 1311–1326. 10.1890/0012-9658(2000)081%5B1311:BDIFNA%5D2.0.CO;2

7. Björk, J. R. (2019). Vertical transmission of sponge microbiota is inconsistent and unfaithful. 3.

8. Boutin, S., Wauters, L. A., McAdam, A. G., Humphries, M. M., Tosi, G., & Dhondt, A. A. (2006). Anticipatory Reproduction and Population Growth in Seed Predators. Science, 314(5807), 1928–1930. 10.1126/science.1135520

9. Bruce-Keller, A. J., Salbaum, J. M., Luo, M., Blanchard, E., Taylor, C. M., Welsh, D. A., & Berthoud, H.-R. (2015). Obese-type Gut Microbiota Induce Neurobehavioral Changes in the Absence of Obesity. Biological Psychiatry, 77(7), 607–615. 10.1016/j.biopsych.2014.07.012

10. Bruijning, M., Henry, L. P., Forsberg, S. K. G., Metcalf, C. J. E., & Ayroles, J. F. (2022). Natural selection for imprecise vertical transmission in host–microbiota systems. 6.

11. Bürkner, P.-C. (2017). brms: An *R* Package for Bayesian Multilevel Models Using *Stan*. Journal of Statistical Software, 80(1). 10.18637/jss.v080.i01

12. Callahan, B. J., McMurdie, P. J., Rosen, M. J., Han, A. W., Johnson, A. J. A., & Holmes, S. P. (2016). DADA2: High-resolution sample inference from Illumina amplicon data. Nature Methods, 13(7), 581–583. 10.1038/nmeth.3869

13. Carey, H. V., & Assadi-Porter, F. M. (2017). The Hibernator Microbiome: Host-Bacterial Interactions in an Extreme Nutritional Symbiosis. Annual Review of Nutrition, 37(1), 477– 500. 10.1146/annurev-nutr-071816-064740

14. Chen, Y., Wang, Y., Shaoyong, W., He, Y., Liu, Y., Wei, S., Gan, Y., Sun, L., Wang, Y., Zong, X., Xiang, Y., Wang, Y., & Jin, M. (2025). High-fertility sows reshape gut microbiota: The rise of serotonin-related bacteria and its impact on sustaining reproductive performance. Journal of Animal Science and Biotechnology, 16(1), 73. 10.1186/s40104-025-01191-z

15. Cheng, C. C., Duar, R. M., Lin, X., Perez-Munoz, M. E., Tollenaar, S., Oh, J.-H., Van Pijkeren, J.-P., Li, F., Van Sinderen, D., Gänzle, M. G., & Walter, J. (2020). Ecological Importance of Cross-Feeding of the Intermediate Metabolite 1,2-Propanediol between Bacterial Gut Symbionts. Applied and Environmental Microbiology, 86(11), e00190–20. 10.1128/AEM.00190-20

16. Clotfelter, E. D., Pedersen, A. B., Cranford, J. A., Ram, N., Snajdr, E. A., Nolan, V., & Ketterson, E. D. (2007). Acorn mast drives long-term dynamics of rodent and songbird populations. Oecologia, 154(3), 493–503. 10.1007/s00442-007-0859-z

17. Dantzer, B., McAdam, A. G., Humphries, M. M., Lane, J. E., & Boutin, S. (2020). Decoupling the effects of food and density on life-history plasticity of wild animals using field experiments: Insights from the steward who sits in the shadow of its tail, the North American red squirrel. Journal of Animal Ecology, 89(11), 2397–2414. 10.1111/1365-2656.13341

18. Downie, B., & Bewley, J. D. (2000). Soluble sugar content of white spruce (*Picea glauca*) seeds during and after germination. Physiologia Plantarum, 110(1), 1–12. 10.1034/j.1399-3054.2000.110101.x

19. Esquivel-Elizondo, S., Ilhan, Z. E., Garcia-Peña, E. I., & Krajmalnik-Brown, R. (2017). Insights into Butyrate Production in a Controlled Fermentation System via Gene Predictions. mSystems, 2(4). 10.1128/msystems.00051-17

20. Fletcher, Q. E., Boutin, S., Lane, J. E., LaMontagne, J. M., McAdam, A. G., Krebs, C. J., & Humphries, M. M. (2010). The functional response of a hoarding seed predator to mast seeding. Ecology, 91(9), 2673–2683. 10.1890/09-1816.1

21. Fletcher, Q. E., Speakman, J. R., Boutin, S., McAdam, A. G., Woods, S. B., & Humphries, M. M. (2012). Seasonal stage differences overwhelm environmental and individual factors as determinants of energy expenditure in free-ranging red squirrels: Cost of living in wild squirrels. Functional Ecology, 26(3), 677–687. 10.1111/j.1365-2435.2012.01975.x

22. Gruninger, R. J., McAllister, T. A., & Forster, R. J. (2016). Bacterial and Archaeal Diversity in the Gastrointestinal Tract of the North American Beaver (Castor canadensis). PLOS ONE, 11(5), e0156457. 10.1371/journal.pone.0156457

23. Hämäläinen, A., McAdam, A. G., Dantzer, B., Lane, J. E., Haines, J. A., Humphries, M. M., & Boutin, S. (2017). Fitness consequences of peak reproductive effort in a resource pulse system. Scientific Reports, 7(1), 9335. 10.1038/s41598-017-09724-x

24. Hartinger, T., Pacífico, C., Poier, G., Terler, G., Klevenhusen, F., & Zebeli, Q. (2022). Shift of dietary carbohydrate source from milk to various solid feeds reshapes the rumen and fecal microbiome in calves. Scientific Reports, 12(1), 12383. 10.1038/s41598-022-16052-2

25. Henry, L. P., Bruijning, M., Forsberg, S. K. G., & Ayroles, J. F. (2021). The microbiome extends host evolutionary potential. Nature Communications, 12(1). 10.1038/s41467-021-25315-x

26. Hicks, A. L., Lee, K. J., Couto-Rodriguez, M., Patel, J., Sinha, R., Guo, C., Olson, S. H., Seimon, A., Seimon, T. A., Ondzie, A. U., Karesh, W. B., Reed, P., Cameron, K. N., Lipkin, W. I., & Williams, B. L. (2018). Gut microbiomes of wild great apes fluctuate seasonally in response to diet. Nature Communications, 9(1), 1786. 10.1038/s41467-018-04204-w

27. Hooper, L. V., Littman, D. R., & Macpherson, A. J. (2012). Interactions Between the Microbiota and the Immune System. Science, 336(6086), 1268–1273. 10.1126/science.1223490

28. Kohl, K. D., Miller, A. W., Marvin, J. E., Mackie, R., & Dearing, M. D. (2014). Herbivorous rodents (Neotoma spp.) harbour abundant and active foregut microbiota. Environmental Microbiology, 16(9), 2869–2878. 10.1111/1462-2920.12376

29. Kolodny, O., & Schulenburg, H. (2020). Microbiome-mediated plasticity directs host evolution along several distinct time scales. Philosophical Transactions of the Royal Society B: Biological Sciences, 375(1808), 20190589. 10.1098/rstb.2019.0589

30. Kubinec, R. (2023). Ordered Beta Regression: A Parsimonious, Well-Fitting Model for Continuous Data with Lower and Upper Bounds. Political Analysis, 31(4), 519–536. 10.1017/pan.2022.20

31. Lach, G., Schellekens, H., Dinan, T. G., & Cryan, J. F. (2018). Anxiety, Depression, and the Microbiome: A Role for Gut Peptides. Neurotherapeutics, 15(1), 36–59. 10.1007/s13311-017-0585-0

32. Langille, M. G. I., Zaneveld, J., Caporaso, J. G., McDonald, D., Knights, D., Reyes, J. A., Clemente, J. C., Burkepile, D. E., Vega Thurber, R. L., Knight, R., Beiko, R. G., & Huttenhower, C. (2013). Predictive functional profiling of microbial communities using 16S rRNA marker gene sequences. Nature Biotechnology, 31(9), 814–821. 10.1038/nbt.2676

33. Leftwich, P. T., Edgington, M. P., & Chapman, T. (2020). Transmission efficiency drives host–microbe associations. Proceedings of the Royal Society B: Biological Sciences, 287(1934), 20200820. 10.1098/rspb.2020.0820

34. Lenth, R., & Piaskowski, J. (2025). *emmeans: Estimated Marginal Means, aka Least-Squares Means* (Version 2.0.1) [Computer software]. https://rvlenth.github.io/emmeans/

35. Levins, R. (1968). Evolution in Changing Environments. Princeton University Press. 10.1515/9780691209418

36. Ma, Z. (Sam). (2020). Testing the Anna Karenina Principle in Human Microbiome-Associated Diseases. iScience, 23(4), 101007. 10.1016/j.isci.2020.101007

37. Marshal, J. P., Owen-Smith, N., Whyte, I. J., & Stenseth, N. Chr. (2011). The role of El Niño–Southern Oscillation in the dynamics of a savanna large herbivore population. Oikos, 120(8), 1175–1182. 10.1111/j.1600-0706.2010.19155.x

38. Maurice, C. F., Knowles, S. C. L., Ladau, J., Pollard, K. S., Fenton, A., Pedersen, A. B., & Turnbaugh, P. J. (2015). Marked seasonal variation in the wild mouse gut microbiota. The ISME Journal, 9(11), 2423–2434. 10.1038/ismej.2015.53

39. McAdam, A. G., Boutin, S., Dantzer, B., & Lane, J. E. (2019). Seed Masting Causes Fluctuations in Optimum Litter Size and Lag Load in a Seed Predator. The American Naturalist, 194(4), 574–589. 10.1086/703743

40. Moeller, A. H., & Sanders, J. G. (2020). Roles of the gut microbiota in the adaptive evolution of mammalian species. Philosophical Transactions of the Royal Society B, 375(1808), 20190597. 10.1098/rstb.2019.0597

41. Moeller, A. H., Suzuki, T. A., Phifer-Rixey, M., & Nachman, M. W. (2018). Transmission modes of the mammalian gut microbiota. Science, 362(6413), 453–457. 10.1126/science.aat7164

42. Moya, A., & Ferrer, M. (2016). Functional Redundancy-Induced Stability of Gut Microbiota Subjected to Disturbance. Trends in Microbiology, 24(5), 402–413. 10.1016/j.tim.2016.02.002

43. Murphy, K. M., Le, S. M., Wilson, A. E., & Warner, D. A. (2023). The Microbiome as a Maternal Effect: A Systematic Review on Vertical Transmission of Microbiota. Integrative And Comparative Biology, 63(3), 597–609. 10.1093/icb/icad031

44. Nienstaedt, H., & Zasada, J. (2008). White Spruce (Picea glauca (Moench) Voss). In Safety Assessment of Transgenic Organisms (pp. 204–236). OECD Publishing. 10.1787/9789264053465-24-en

45. Oksanen, J., Blanchet, F. G., Friendly, M., Kindt, R., Legendre, P., McGlinn, D., Minchin, P. R., O’Hara, R. B., Simpson, G. L., Solymos, P., Stevens, M. H. H., Szoecs, E., & Wagner, H. (2018). *vegan: Community Ecology Package*. R package version, 2–5. https://cran.r-project.org/package=vegan

46. Petrullo, L., Boutin, S., Lane, J. E., McAdam, A. G., & Dantzer, B. (2023). Phenotype-environment mismatch errors enhance lifetime fitness in wild red squirrels. Science, 379(6629), 269–272. 10.1126/science.abn0665

47. Petrullo, L., Morris, N. J., Tharin, C., & Dantzer, B. (2025). Harbingers of change: Towards a mechanistic understanding of anticipatory plasticity in animal systems. Functional Ecology, 39(11), 2999–3020. 10.1111/1365-2435.70071

48. Petrullo, L., Ren, T., Wu, M., Boonstra, R., Palme, R., Boutin, S., McAdam, A. G., & Dantzer, B. (2022). Glucocorticoids coordinate changes in gut microbiome composition in wild North American red squirrels. Scientific Reports, 12(1), 1–12. 10.1038/s41598-022-06359-5

49. Petrullo, L., Webber, Q., Raulo, A., Boutin, S., Lane, J. E., McAdam, A. G., & Dantzer, B. (2025). Social Microbial Transmission in a Solitary Mammal. Ecology Letters, 28(8), e70186. 10.1111/ele.70186

50. Pullman, G. S., Chase, K.-M., Skryabina, A., & Bucalo, K. (2008). Conifer embryogenic tissue initiation: Improvements by supplementation of medium with D-xylose and D-chiro-inositol. Tree Physiology, 29(1), 147–156. 10.1093/treephys/tpn013

51. Quast, C., Pruesse, E., Yilmaz, P., Gerken, J., Schweer, T., Yarza, P., Peplies, J., & Glöckner, F. O. (2012). The SILVA ribosomal RNA gene database project: Improved data processing and web-based tools. Nucleic Acids Research, 41(D1), 590–596. 10.1093/nar/gks1219

52. Raulo, A., Bürkner, P. C., Finerty, G. E., Dale, J., Hanski, E., English, H. M., Lamberth, C., Firth, J. A., Coulson, T., & Knowles, S. C. L. (2024). Social and environmental transmission spread different sets of gut microbes in wild mice. Nature Ecology and Evolution, 8(5), 972–985. 10.1038/s41559-024-02381-0

53. Ren, T., Boutin, S., Humphries, M. M., Dantzer, B., Gorrell, J. C., Coltman, D. W., McAdam, A. G., & Wu, M. (2017). Seasonal, spatial, and maternal effects on gut microbiome in wild red squirrels. Microbiome, 5(1), 163. 10.1186/s40168-017-0382-3

54. Rusling, M., Karim, A., Kaye, A., Lee, C.-M. J., Wegman-Points, L., Mathis, V., Lampeter, T., & Yuan, L.-L. (2024). Influences of Ruminococcus bromii and Peptostreptococcaceae on voluntary exercise behavior in a rodent model. Frontiers in Microbiomes, 3, 1389103. 10.3389/frmbi.2024.1389103

55. Stothart, M. R., Lavergne, S., McCaw, L., Singh, H., De Vega, W., Amato, K., Poissant, J., & Boonstra, R. (2025). Population Dynamics and the Microbiome in a Wild Boreal Mammal: The Snowshoe Hare Cycle and Impacts of Diet, Season and Predation Risk. Molecular Ecology, 34(3), e17629. 10.1111/mec.17629

56. Taylor, R. W., Boon, A. K., Dantzer, B., Réale, D., Humphries, M. M., Boutin, S., Gorrell, J. C., Coltman, D. W., & McADAM, A. G. (2012). Low heritabilities, but genetic and maternal correlations between red squirrel behaviours. Journal of Evolutionary Biology, 25(4), 614–624. 10.1111/j.1420-9101.2012.02456.x

57. Team R Development Core. (2020). A Language and Environment for Statistical Computing. In R Foundation for Statistical Computing. R Foundation for Statistical Computing. http://www.r-project.org

58. Valles-Colomer, M., Blanco-Míguez, A., Manghi, P., Asnicar, F., Dubois, L., Golzato, D., Armanini, F., Cumbo, F., Huang, K. D., Manara, S., Masetti, G., Pinto, F., Piperni, E., Punčochář, M., Ricci, L., Zolfo, M., Farrant, O., Goncalves, A., Selma-Royo, M., … Segata, N. (2023). The person-to-person transmission landscape of the gut and oral microbiomes. Nature, 614(7946), 125–135. 10.1038/s41586-022-05620-1

59. Varpe, Ø. (2017). Life History Adaptations to Seasonality. Integrative and Comparative Biology, 57(5), 943–960. 10.1093/icb/icx123

60. Wang, Q., Garrity, G. M., Tiedje, J. M., & Cole, J. R. (2007). Naïve Bayesian classifier for rapid assignment of rRNA sequences into the new bacterial taxonomy. Applied and Environmental Microbiology, 73(16), 5261–5267. 10.1128/AEM.00062-07

61. Webber, Q. M. R., Dantzer, B., Lane, J. E., Boutin, S., & McAdam, A. G. (2023). Density-dependent plasticity in territoriality revealed using social network analysis. Journal of Animal Ecology, 92(1), 207–221. 10.1111/1365-2656.13846

62. Week, B., Russell, S. L., Schulenburg, H., Bohannan, B. J. M., & Bruijning, M. (2025). Applying evolutionary theory to understand host–microbiome evolution. Nature Ecology & Evolution, 9(10), 1769–1780. 10.1038/s41559-025-02846-w

63. Yang, L. H. (2004). Periodical Cicadas as Resource Pulses in North American Forests. Science, 306(5701), 1565–1567. 10.1126/science.1103114

64. Zaneveld, J. R., McMinds, R., & Vega Thurber, R. (2017). Stress and stability: Applying the Anna Karenina principle to animal microbiomes. Nature Microbiology, 2(9), 17121. 10.1038/nmicrobiol.2017.121

65. Zhu, H., Li, G., Liu, J., Xu, X., & Zhang, Z. (2022). Gut microbiota is associated with the effect of photoperiod on seasonal breeding in male Brandt’s voles (Lasiopodomys brandtii). Microbiome, 10(1), 194. 10.1186/s40168-022-01381-1

