## Supplementary information for "Maternal effects shape scale-dependent convergence in gut microbial responses to environmental change"

**Supplementary information: Maternal effects disrupt scale-dependent divergence in gut microbial responses to environmental change**

**Authors:** Gabriele Schiro^1,2^, Allyson Placko^1^, Stan Boutin^3^, Ben Dantzer^4,5^, Jeffery Lane^6^, Andrew G. McAdam^7^, and Lauren Petrullo^1^

**Supplementary information S1**.

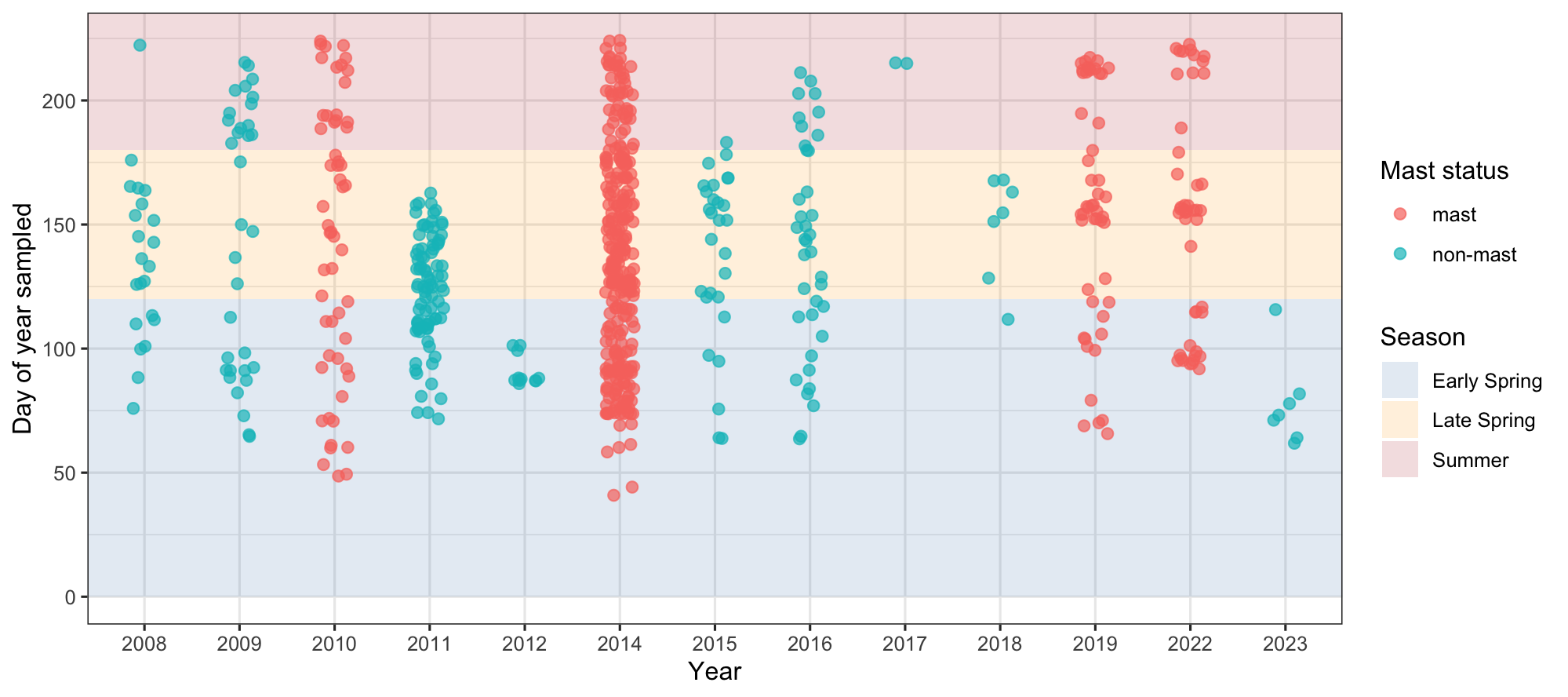

**Supplementary information S1**. Seasonal timing of observations by year and mast status. Points show the day of year each sample was collected from 2008–2023, colored by mast (pink) and non-mast (teal) years. Background shading indicates seasonal periods: Early Spring (days 0–120), Late Spring (121–180), and Summer (>180).

**Supplementary information S2**.

Output of a linear mixed-effects model testing effects of season, mast status, and their interaction on alpha diversity (richness), controlling for age, sex, group, and read number. Random intercepts were included for individual (squirrel ID), year, and sequencing run. Estimates, 95% confidence intervals, and p-values are shown.

Formula:

*Richness ~ season_cat * mast + age + sex + gr + read_number + (1 | squirrel_id_letter) + (1 | year_factor) + (1 | run.x).*

|  | **Richness** | | |
| --- | --- | --- | --- |
| *Predictors* | *Estimates* | *CI* | *p* |
| (Intercept) | 200.26 | 140.50 – 260.01 | **<0.001** |
| season cat [Late Spring] | -30.29 | -38.54 – -22.05 | **<0.001** |
| season cat [Summer] | -9.31 | -19.14 – 0.53 | 0.064 |
| mast [non-mast] | -13.14 | -27.08 – 0.80 | 0.065 |
| age | -2.63 | -4.80 – -0.46 | **0.018** |
| sex [M] | -5.73 | -20.12 – 8.65 | 0.434 |
| gr [SU] | -3.50 | -15.88 – 8.88 | 0.579 |
| read number | 0.00 | 0.00 – 0.00 | **<0.001** |
| season cat [Late Spring]  × mast [non-mast] | 25.14 | 11.74 – 38.54 | **<0.001** |
| season cat [Summer] ×  mast [non-mast] | -33.31 | -52.45 – -14.17 | **0.001** |
| **Random Effects** | | | |
| σ^2^ | 1322.88 | | |
| τ_00_ _squirrel_id_letter_ | 649.04 | | |
| τ_00_ _year_factor_ | 0.00 | | |
| τ_00_ _run.x_ | 2483.54 | | |
| N _squirrel_id_letter_ | 112 | | |
| N _year_factor_ | 13 | | |
| N _run.x_ | 3 | | |
| Observations | 679 | | |
| Marginal R^2^ / Conditional R^2^ | 0.889 / NA | | |

**Supplementary information S3**.

Output of a linear mixed-effects model testing effects of season, mast status, and their interaction on Tukey-transformed alpha diversity (Shannon H’), controlling for age, sex, group, and read number. Random intercepts were included for individual (squirrel ID), year, and sequencing run. Estimates, 95% confidence intervals, and p-values are shown.

Formula:

*Tukey transformed Shannon H’ ~ season_cat * mast + age + sex + gr + read_number + (1 | squirrel_id_letter) + (1 | year_factor) + (1 | run.x).*

|  | **Tukey shannon** | | |
| --- | --- | --- | --- |
| *Predictors* | *Estimates* | *CI* | *p* |
| (Intercept) | 5853.99 | 3999.35 – 7708.62 | **<0.001** |
| season cat [Late Spring] | -909.69 | -1271.65 – -547.73 | **<0.001** |
| season cat [Summer] | -1019.87 | -1449.93 – -589.81 | **<0.001** |
| mast [non-mast] | -90.28 | -986.30 – 805.75 | 0.843 |
| age | -61.07 | -169.44 – 47.30 | 0.269 |
| sex [M] | -96.02 | -618.38 – 426.34 | 0.718 |
| gr [SU] | -168.17 | -609.32 – 272.97 | 0.454 |
| read number | 0.04 | 0.03 – 0.05 | **<0.001** |
| season cat [Late Spring]  × mast [non-mast] | 597.04 | -6.15 – 1200.22 | 0.052 |
| season cat [Summer] ×  mast [non-mast] | -1098.18 | -1983.84 – -212.52 | **0.015** |
| **Random Effects** | | | |
| σ^2^ | 2581016.10 | | |
| τ_00_ _squirrel_id_letter_ | 562230.47 | | |
| τ_00_ _year_factor_ | 281946.49 | | |
| τ_00_ _run.x_ | 1935064.21 | | |
| ICC | 0.52 | | |
| N _squirrel_id_letter_ | 112 | | |
| N _year_factor_ | 13 | | |
| N _run.x_ | 3 | | |
| Observations | 679 | | |
| Marginal R^2^ / Conditional R^2^ | 0.231 / 0.630 | | |

**Supplementary information S4.** Full results of PERMANOVA testing the effects of sequencing run, host traits, and environmental variables on microbial community composition. Analyses were performed using 999 permutations with strata set to individual identity (squirrel ID) to account for repeated measures. Terms were added sequentially (type I sums of squares; *by* = "terms"). R² values represent the proportion of variance explained by each predictor. Significant p-values (≤ 0.05) are indicated in bold.

| **Predictor** | **Df** | **Sum of Squares** | **R²** | **F** | **p-value** |
| --- | --- | --- | --- | --- | --- |
| run | 2 | 9.760 | 0.0556 | 23.44 | 0.001 |
| sex | 1 | 0.987 | 0.0056 | 4.74 | 0.001 |
| gr | 1 | 1.365 | 0.0078 | 6.56 | 0.001 |
| age | 1 | 0.926 | 0.0053 | 4.45 | 0.001 |
| year_factor | 12 | 15.037 | 0.0857 | 6.02 | 0.001 |
| season_cat | 2 | 9.009 | 0.0514 | 21.64 | 0.001 |
| season_cat × mast | 2 | 1.346 | 0.0077 | 3.23 | 0.001 |
| Residual | 658 | 136.982 | 0.7809 | — | — |
| **Total** | 679 | 175.412 | 1.0000 | — | — |

**Supplementary information S5**

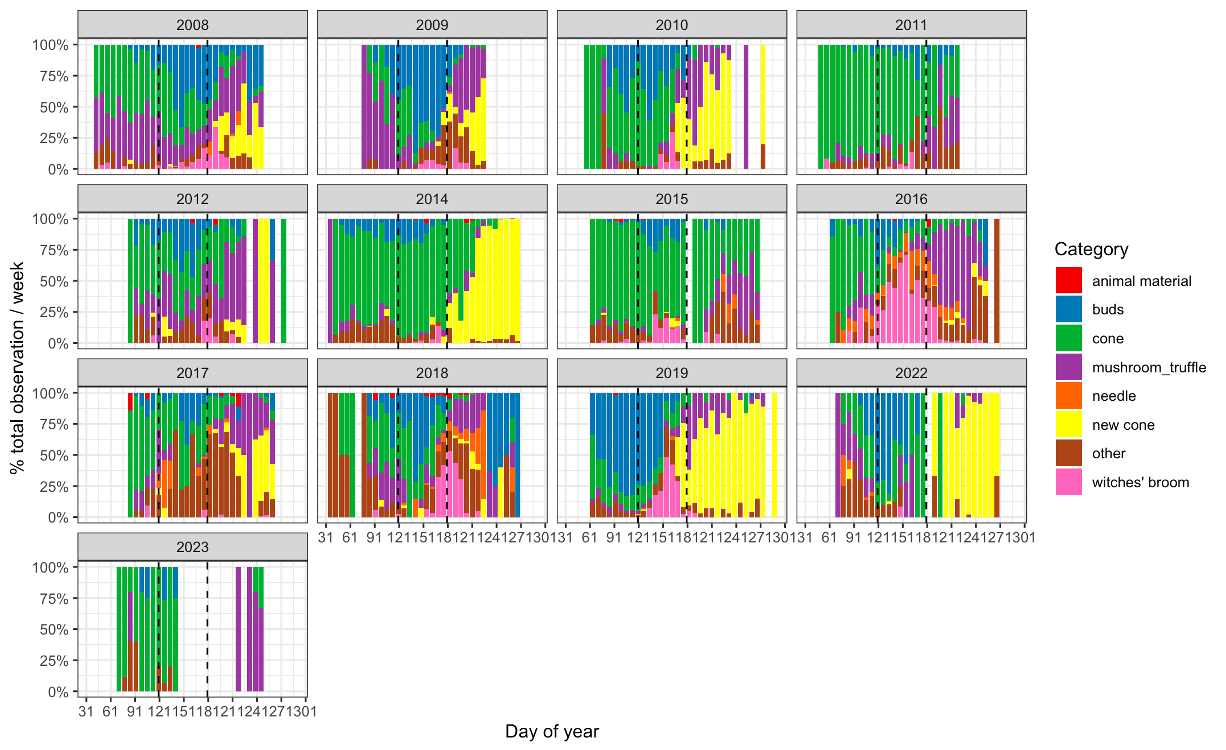

**Supplementary information S5.** Seasonal and interannual variation in squirrel diet composition and its relationship with gut microbiome turnover. Stacked bar plots show weekly relative dietary composition across years based on observational feeding categories. Bars represent the proportional contribution of each food category within a given week, and dashed vertical lines indicate selected seasonal reference points within each year.

**Supplementary information S8**
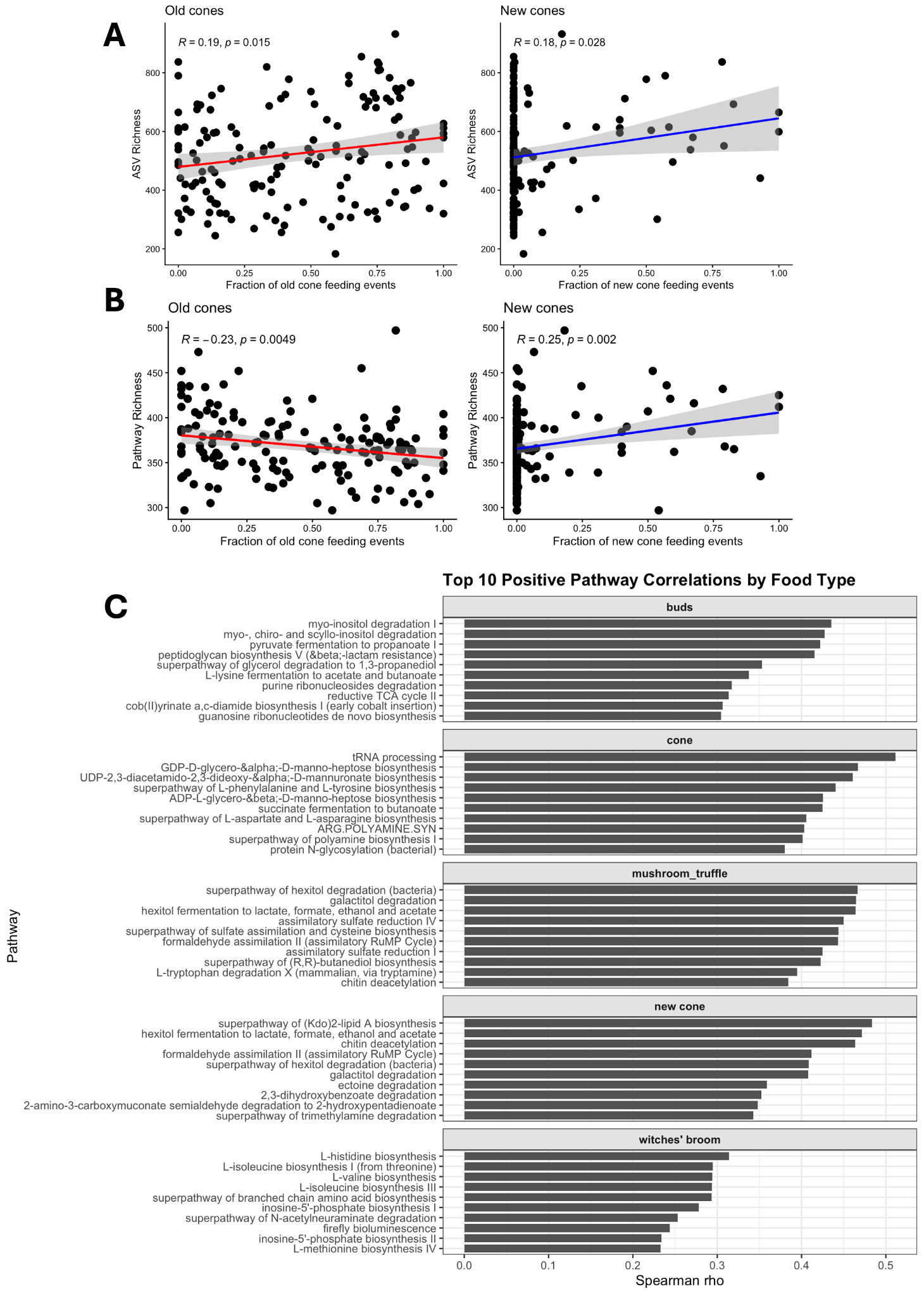

**Supplementary information S8. Associations between cone-feeding behavior and microbial diversity and function.** Top panels show relationships between the fraction of feeding events on old cones or new cones and bacterial ASV richness (top row) or inferred metabolic pathway richness (bottom row). Lines represent linear regressions with shaded 95% confidence intervals; Pearson correlation coefficients (*R*) and associated *p*-values are shown within each panel. Feeding on old cones was associated with increased taxonomic richness but reduced functional richness, consistent with greater functional redundancy among microbial taxa. In contrast, feeding on new cones was positively associated with both taxonomic and functional richness, suggesting broader metabolic functional enrichment and increased functional diversification.

Bottom panels show the top 10 positively correlated microbial metabolic pathways for each food type based on Spearman’s rho. Bud-associated pathways were dominated by inositol/polyol degradation and fermentation pathways, indicating a focused carbohydrate-utilization signature, with specialization in conifer buds as inositol is found in conifer buds. Old cone-associated pathways were enriched for nucleotide biosynthesis, tRNA processing,and various biosytehsis of polysaccardies and cell components indicating, possibly indicating specialized degradation of such compounds (often biosynthesis and degradation pathways share genetic machinery). Mushroom/truffle-associated pathways were enriched for sulfur assimilation, formaldehyde metabolism and chitin processing pathways, suggesting utilization of fungal-derived substrates. In contrast, new cone-associated pathways spanned diverse metabolic classes, including lipid biosynthesis, fermentation, chitin metabolism, osmolyte degradation, and aromatic compound degradation, consistent with broader functional diversification. Witches’ broom samples were enriched primarily for amino acid and other biosynthetic pathways, indicating an unknown possibly specialized anabolic functional profile.

**Supplementary information S9.** BRMS full model setting and outputs, Bray-Curtis similarity. Model output is in original output model scale (logit scale) before conversion to response scale.

Model Specification Bray Curtis model

*Family: beta*

*Links: mu = logit; phi = identity*

*Formula:*

*BC_Similarity_rar ~ 1 + mother_offspring * masting +*

*mother_offspring * same_season + year_dist + same_year +*

*dist_m + same_sex + run + age_dist +*

*(1 | mm(Sample1, Sample2)) + (1 | mm(squirrel1, squirrel2))*

*Data: melted_diffs_selected_try (N = 262,450 observations)*

*Draws: 4 chains x 3000 iter (1000 warmup); 8000 post-warmup draws*

*Multilevel Hyperparameters*

*~mmSample1Sample2 (Number of levels: 725)*

| Parameter | Estimate | Est.Error | l-95% CI | u-95% CI | Rhat | Bulk_ESS | Tail_ESS |
| --- | --- | --- | --- | --- | --- | --- | --- |
| sd(Intercept) | 0.38 | 0.01 | 0.36 | 0.40 | 1.00 | 1397 | 3263 |

~mmsquirrel1squirrel2 (Number of levels: 110)

| Parameter | Estimate | Est.Error | l-95% CI | u-95% CI | Rhat | Bulk_ESS | Tail_ESS |
| --- | --- | --- | --- | --- | --- | --- | --- |
| sd(Intercept) | 0.12 | 0.03 | 0.06 | 0.17 | 1.02 | 258 | 213 |

Regression Coefficients

| Parameter | Estimate | Est.Error | l-95% CI | u-95% CI | Rhat | Bulk_ESS | Tail_ESS |
| --- | --- | --- | --- | --- | --- | --- | --- |
| Intercept | -0.91 | 0.02 | -0.96 | -0.88 | 1.00 | 2850 | 3948 |
| mother_offspring | 0.02 | 0.01 | 0.00 | 0.04 | 1.00 | 7918 | 5141 |
| masting | -0.06 | 0.00 | -0.07 | -0.06 | 1.00 | 6878 | 5570 |
| same_season | 0.18 | 0.00 | 0.18 | 0.18 | 1.00 | 8497 | 5471 |
| year_dist | -0.26 | 0.00 | -0.27 | -0.25 | 1.00 | 7042 | 5721 |
| same_year | 0.12 | 0.00 | 0.11 | 0.12 | 1.00 | 7559 | 6820 |
| dist_m | -0.11 | 0.00 | -0.12 | -0.10 | 1.00 | 6923 | 4289 |
| same_sex | 0.01 | 0.00 | 0.00 | 0.01 | 1.00 | 8003 | 4905 |
| run | 0.05 | 0.00 | 0.04 | 0.05 | 1.00 | 7395 | 6780 |
| age_dist | -0.01 | 0.00 | -0.01 | -0.00 | 1.00 | 7867 | 5235 |
| mother_offspring:masting | 0.02 | 0.01 | 0.00 | 0.04 | 1.00 | 6836 | 5245 |
| mother_offspring:same_season | 0.01 | 0.01 | -0.02 | 0.03 | 1.00 | 7727 | 5834 |

Further Distributional Parameters

| Parameter | Estimate | Est.Error | l-95% CI | u-95% CI | Rhat | Bulk_ESS | Tail_ESS |
| --- | --- | --- | --- | --- | --- | --- | --- |
| phi | 84.82 | 0.23 | 84.35 | 85.28 | 1.00 | 7246 | 4717 |

Draws were sampled using sample(hmc). Bulk_ESS and Tail_ESS are effective sample size measures. Rhat is the potential scale reduction factor on split chains (at convergence, Rhat = 1).

**Supplementary information S10.** BRMS full model setting and outputs, Jaccard similarity. Model output is in the original output model scale (logit scale) before conversion to response scale.

Model Specification Jaccard

*Family: beta*

*Links: mu = logit; phi = identity*

*Formula:*

*Jaccard_Similarity ~ 1 + mother_offspring * masting +*

*mother_offspring * same_season + same_year + year_dist +*

*dist_m + same_sex + read_dist + run + age_dist +*

*(1 | mm(Sample1, Sample2)) + (1 | mm(squirrel1, squirrel2))*

*Data: melted_diffs_selected_try (N = 262,450 observations)*

*Draws: 4 chains x 3000 iter (1000 warmup); 8000 post-warmup draws*

*Multilevel Hyperparameters*

*~mmSample1Sample2 (Number of levels: 725)*

| Parameter | Estimate | Est.Error | l-95% CI | u-95% CI | Rhat | Bulk_ESS | Tail_ESS |
| --- | --- | --- | --- | --- | --- | --- | --- |
| sd(Intercept) | 0.25 | 0.01 | 0.23 | 0.26 | 1.00 | 3471 | 5321 |

~mmsquirrel1squirrel2 (Number of levels: 110)

| Parameter | Estimate | Est.Error | l-95% CI | u-95% CI | Rhat | Bulk_ESS | Tail_ESS |
| --- | --- | --- | --- | --- | --- | --- | --- |
| sd(Intercept) | 0.24 | 0.02 | 0.20 | 0.29 | 1.00 | 3411 | 5472 |

Regression Coefficients

| Parameter | Estimate | Est.Error | l-95% CI | u-95% CI | Rhat | Bulk_ESS | Tail_ESS |
| --- | --- | --- | --- | --- | --- | --- | --- |
| Intercept | -0.80 | 0.03 | -0.86 | -0.75 | 1.00 | 10751 | 6160 |
| mother_offspring | 0.02 | 0.01 | 0.01 | 0.04 | 1.00 | 8904 | 5383 |
| masting | 0.03 | 0.00 | 0.02 | 0.03 | 1.00 | 8021 | 5441 |
| same_season | 0.13 | 0.00 | 0.12 | 0.13 | 1.00 | 8166 | 6872 |
| same_year | 0.10 | 0.00 | 0.10 | 0.10 | 1.00 | 7931 | 6930 |
| year_dist | -0.01 | 0.00 | -0.01 | -0.00 | 1.00 | 8156 | 5835 |
| dist_m | -0.09 | 0.00 | -0.10 | -0.09 | 1.00 | 8921 | 5026 |
| same_sex | 0.01 | 0.00 | 0.01 | 0.01 | 1.00 | 7653 | 4642 |
| read_dist | -0.49 | 0.00 | -0.50 | -0.49 | 1.00 | 8156 | 5450 |
| run | -0.01 | 0.00 | -0.02 | -0.01 | 1.00 | 7643 | 7447 |
| age_dist | -0.03 | 0.00 | -0.03 | -0.03 | 1.00 | 7991 | 5264 |
| mother_offspring:masting | 0.02 | 0.01 | 0.00 | 0.03 | 1.00 | 8423 | 5873 |
| mother_offspring:same_season | 0.01 | 0.01 | -0.00 | 0.03 | 1.00 | 8611 | 6217 |

Further Distributional Parameters

| Parameter | Estimate | Est.Error | l-95% CI | u-95% CI | Rhat | Bulk_ESS | Tail_ESS |
| --- | --- | --- | --- | --- | --- | --- | --- |
| phi | 181.65 | 0.51 | 180.66 | 182.64 | 1.00 | 8580 | 4569 |

Draws were sampled using sample(hmc). Bulk_ESS and Tail_ESS are effective sample size measures. Rhat is the potential scale reduction factor on split chains (at convergence, Rhat = 1).
